# Novel frontier in wildlife monitoring: identification of small rodent species from faecal pellets using Near-Infrared Reflectance Spectroscopy (NIRS)

**DOI:** 10.1101/2022.04.11.487830

**Authors:** Maria W. Tuomi, Francisco J.A. Murguzur, Katrine S. Hoset, Eeva M. Soininen, Eero Vesterinen, Tove Aa. Utsi, Sissel Kaino, Kari Anne Bråthen

## Abstract

Small rodents are prevalent and functionally important across world’s biomes, making their monitoring salient for ecosystem management, conservation, forestry and agriculture. Yet, there is a dearth of cost-effective and non-invasive methods for large-scale, intensive sampling. As one such method, fecal pellet counts readily provide relative abundance indices. Given available analytical methods, feces could also allow for determination of multiple ecological and physiological variables, including community composition. We developed calibration models for rodent taxonomic determination using fecal near-infrared reflectance spectroscopy (fNIRS).

Our results demonstrate fNIRS as an accurate and robust method for predicting genus and species identity of five co-existing subarctic microtine rodent species. We show that sample exposure to weathering did not reduce accuracy, indicating suitability of the method for samples collected from the field. Diet was not a major determinant of species prediction accuracy in our samples, as diet exhibited large variation and overlap between species. While regional calibration models predicted poorly samples from another region, calibration models including samples from two regions provided a good prediction accuracy for both regions.

We propose fNIRS as a fast and cost-efficient high-throughoutput method for rodent taxonomic determination, and highlight its potential for cross-regional calibrations and use on field-collected samples. FNIRS can facilitate rodent population censuses at larger spatial extent than before deemed feasible, if combined with pellet-count based abundance indices. Given the versatility of fNIRS analytics, developing such monitoring schemes can support ecosystem- and interaction-based approaches to monitoring.

## INTRODUCTION

Small rodents are prevalent in ecosystems across the globe, with many species acting as ecosystem engineers (Dickman, 1999) or keystone species (Ims & Fuglei, 2005); with numerous rare species of high conservation value (Green, Ting, Manjerovic, & Mateus-Pinilla, 2013) and many invasive populations with profound effects on ecosystem functioning (Dickman, 1999). Monitoring and research of small rodents is thus a globally salient enterprise for ecosystem management, conservation, forestry and agriculture. Yet, obtaining population- or community-level data on small rodents is often challenging, as these small and cryptic animals are elusive (Green et al., 2013) and cost-effective methods for large-scale sampling of their multispecies communities are largely missing (Engeman & Whisson, 2006; Heisler, Somers, & Poulin, 2016). State-of-the-art estimation of small rodent community composition, population size and density, relies on a variety of trapping and indexing efforts, e,g, pitfalls, live-, snap-, hair- or camera traps and systematic incidental observations (Engeman & Whisson, 2006; Soininen, Jensvoll, Killengreen, & Ims, 2015; Fauteux et al., 2018), or on counts of burrows, runways, winter nests, owl pellet contents and faeces (Green et al., 2013; Heisler et al., 2016). Trapping methods, while providing actual estimates on species identity, abundance and key demographic parameters, are laboursome (Engeman & Whisson, 2006; Villette, Krebs, Jung, & Boonstra, 2016) and often inadequate for high-intensity sampling with large spatial or temporal coverage (Heisler et al., 2016; Villette et al., 2016).

Multitude of wildlife censuses rely on feces counts (Kohn & Wayne, 1997; Campbell, Swanson, & Sales, 2004; Karels, Koppel, & Hik, 2004; Green et al., 2013). Feces are organic material whose structural and chemical properties are determined by the mechanical, biochemical and microbiological processing of ingested biomass throughout the digestive pathway. Feces could therefore allow for determination of diverse ecological and physiological variables indicative of species interactions (c.f. Ehrich et al., 2019). In addition, feces are readily available samples. Acquiring them does not require being in contact with the animal, thus decreasing risk of infections and animal stress, and feces can provide information from seasons and times when researchers are not present (Kohn & Wayne, 1997; Whisson, Engeman, & Collins, 2005). However, feces counts and other survey-based indexing methods using activity indicators have often limited ability to readily provide information on species identity or demographic parameters (but see Soininen, Gauthier, et al., 2015), unless combined with molecular and endocrinological methods (Schwarzenberger, 2007; Galan, Pagès, & Cosson, 2012). These sample-destructive methods are often too expensive for large sample sizes typical for landscape scale monitoring. Thus, realizing the potential of fecal indices for high-intensity or large-scale sampling calls for more cost-effective high-througoutput analytical methods. Importantly, such methods, if embedded in appropriate monitoring schemes (Yoccoz, Nichols, & Boulinier, 2001) could provide access to questions pertinent to ongoing rapid range shifts and ecosystem changes (Wintle, Runge, & Bekessy, 2010; Lenoir & Svenning, 2015; Pecl et al., 2017). To be feasible and applicable, the method should be robust and sensitive across space and time, utilize easily obtainable samples, be quick in the pre-processing of the samples and be cheap and fast (Whisson et al., 2005; Engeman & Whisson, 2006). Analytical methods that are non-destructive for the fecal samples increase their utility by allowing subsequent application of molecular methods on the samples.

All of these requirements can be met by application of near infrared reflectance spectroscopy (NIRS). NIRS is a widely applied analytic method in agriculture and petrochemical industry (Pasquini, 2003), with increasing representation in different fields of ecology (Villamuelas et al., 2017; Aw & Ballard, 2019; Murguzur et al., 2019). NIRS is a rapid and non-destructive spectral analytic tool for assessing quantitative and qualitative constituents of a wide array of organic and inorganic samples, which requires minimal sample pre-processing, mainly involving sample material homogenization and drying (Pasquini, 2003).

The scanning procedure of a sample takes only seconds, and does not require specialist personnel. Once a calibration model is validated, scanning and analysis of further samples comes with no extra cost of e.g. reagents or genetic primers. Unlike with molecular methods, after NIR-scanning samples can be reused and new calibration models (incl. improved calibrations and modelling new constituents) can be applied to existing spectra. Consequently, fecal NIRS (fNIRS) has been used to predict species identity, demographic parameters (Tolleson, Randel, Stuth, & Neuendorff, 2005; Wiedower et al., 2012; Aw & Ballard, 2019) and diet quality (Foley et al., 1998; Villamuelas et al., 2017) of several wild animals from large mammals to insects. However, the use of fNIRS in wildlife research and monitoring remains largely unknown to the ecologist community, contrasting widespread use of remote sensing (Kerr & Ostrovsky, 2003; Pettorelli et al., 2015) and high-throughoutput genetic barcoding (Yoccoz, 2012) methods.

As with any analytical method, there are potential caveats to using fNIRS. First, as feces are collected from the field they are exposed to ambient weather. This can compromise prediction accuracy because leaching and microbiological processes can change pellet chemical composition (Jenks, Soper, Lochmiller, & Leslie, 1990; Kamler, Homolka, & Kráčmar, 2003). Second, variation between animal individuals may affect their fecal composition and NIR-spectra, and hence decrease or confound prediction accuracy of species. In particular, diet quality affects fNIR-spectra (Stuth, Jama, & Tolleson, 2003; Villamuelas et al., 2017), and may compromise prediction accuracy whenever considerable dietary overlap between species occurs. Especially for coexisting and competing generalist rodent species the effect of diet is likely to be complex, as diets overlap and are dependent on e.g. season and forage item availability (Soininen et al., 2013). In addition, sex and reproductive status may affect fNIR-spectra (Tolleson et al., 2005), increasing variation in the spectral data. Third, if factors driving the species-signal in fNIR-spectra differ between populations, fNIRS calibrations that cover only spatially limited target populations can yield high misclassification rates on samples from a different population. However, recent studies presented NIRS-calibrations applicable across species or geographic regions (Villamuelas et al., 2017; Murguzur et al., 2019), revealing the potential for cross-regional or global NIRS- and fNIRS-calibrations.

Here we develop fNIRS calibration models for rodent taxonomic determination from their pellets (>0.025g of dry weight). Our main hypothesis is that fecal properties differ between taxa, allowing for classification of individuals to genus and species based on their fNIR-spectra (H1). However, this separation capability might erode with the effect of exposure (H2) due to leaching, irradiation and decomposition. Prediction accuracy of individual samples may also be linked with diet composition (H3), which we tested by comparing fNIRS data to DNA-metabarcoding data of the same pellets. Additionally, spectra may display regional differences between populations (H4). For the latter we contrast two sub-hypotheses: H4.1 that calibration models based on samples from one region may perform poorly with samples from other region, and H4.2 that a calibration model including all regions successfully classifies independent test individuals from all regions. Finally, we use our dataset to demonstrate how our calibration model and the modelling framework could be used in practice to predict species identity of fecal samples.

## MATERIAL AND METHODS

We outline the workflow of building NIRS calibrations and testing our hypothesis in Fig. 1. We detail the following steps: rodent fecal sample extraction, sample processing and experimental treatment, NIRS scanning and spectra pre-treatment, calibration modelling including model building, validation and testing, diet molecular analysis including DNA-metabarcoding, bioinformatics and modelling potential confounding of diet, as well as regression modelling of NIRS-calibration model results. Further details on molecular methods and calibration modelling are provided in the Supplementary material.

**Figure 1.**
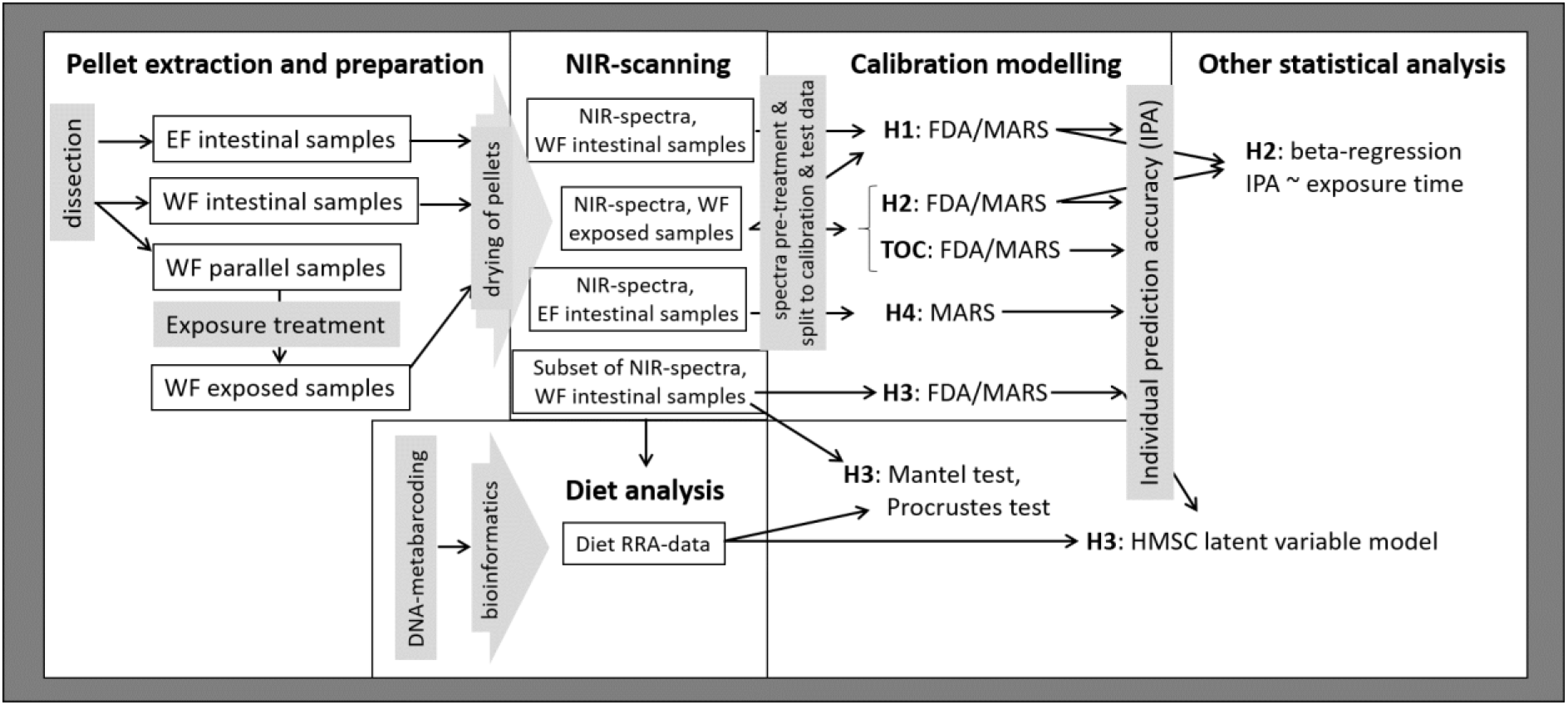
Illustration of the analytical workflow. Detailed description of calibration models is presented in Table 1.

**Table 1.** A summary of calibration model methdology. Column « **Hypothesis »** refers to the the tested question. Column « **Model»** refers to the used model type, i.e. FDA models applying MARS (FDA/MARS) and direct application of MARS. Degree indicates 1-way (degree = 2) or 2-way (degree = 3) interactions of model variables. Number of models indicates that each response variable, e.g. genus or species, has its own calibration model. Column « **Response variable »** describes the categories predicted by the calibration model. Column « **Calibration data»** and « **Test data** » describe the datasets used to calibrate the model and test its generalization error. Total number of rdMCCV iterations per each model is 600, i.e. the product of number of MCCV iterations (n = 200) and number of data splits (n = 3). Abbreviations: WF – West-Finnmark; EF – East-Finnmark; Llem – *Lemmus lemmus*; Mruf – *Myodes rufocanus*; Mrut – *M. rutilus*; Moec – *Microtus oeconomus*; Magr – *M. agrestis*.

| hypothesis | model | response variable | calibration data | test data |
| --- | --- | --- | --- | --- |
| *H1* | FDA/MARS, degree = 2<br>no. of models = 2 | **genus** (Lemmus, Myodes, Microtus); **species** (Llem, Mruf, Mrut, Moec, Magr) | WF intestinal and exposed samples<sup>1</sup> | WF intestinal and exposed samples<sup>1</sup> |
| *H2* | FDA/MARS, degree = 3<br>no. of models = 2 | **genus** (Lemmus, Myodes, Microtus); **species** (Llem, Mruf, Moec, Magr) | WF exposed samples<sup>1</sup> | WF exposed samples<sup>1</sup> |
| *H3* | FDA/MARS, degree = 3<br>no. of models = 1 | **species** (Llem, Mruf, Mrut, Moec, Magr) | WF intestinal samples<sup>2</sup> | WF intestinal samples<sup>1</sup> |
| *H4* | MARS,<br>degree = 3<br>no. of models = 3 | **Mruf, Moec** | H4.1a = WF<sup>1</sup><br>H4.1b = EF<sup>2</sup><br>H4.2 = WF & EF<sup>3</sup> | H4.1a = 100% EF, WF<sup>1</sup><br>H4.1b = 100% WF, EF<sup>2</sup><br>H4.2 = WF & EF<sup>3</sup> |

<sup>1</sup> 95%/5%, 90%/10% and 80%/20% of each species and exposure week
<sup>2</sup> 95%/5%, 90%/10% and 80%/20% of each species
<sup>3</sup> 95%/5%, 90%/10% and 80%/20% of each species, exposure week and region

### Rodent data and fecal samples

We collected fecal samples from animals trapped as part of a long-term rodent population monitoring time-series. The set of rodent individuals included in the calibration modelling incorporated substantial variation in environmental conditions of their trapping site, season and year, as well as variation in individual physiological condition related to reproductive status and age (Supplementary Table S1a). The largest set of individual fecal samples (n=472) came from West-Finnmark, Northern Norway, encompassing two research areas ca. 55km apart (hereafter West-Finnmark or “WF”) – Joatka research area (JRA), (69°45’N, 23°55’E) and Skillefjordnes (Skirvinjárga) (77°86’N, 05°86’E). Small rodent guild at WF includes five out of seven small rodent species occurring throughout Northern Fennoscandia: Norwegian lemmings (*Lemmus lemmus*), grey-sided voles (*Myodes rufocanus*), red voles (*Myodes rutilus*), root voles (*Microtus oeconomus*) and field voles (*Microtus agrestis*). JRA has in total 77 trapping quadrats of 15 by 15m, including snowbed, heath, meadow and wetland habitats. Trapping was conducted bi-annually, during early (mid-June) and late (early-mid September) growing season. Samples were included from a full population cycle between 2010 and 2014, with the largest rodent peak in 40yr occurring in 2011-2012 (see Ekerholm, Oksanen, & Oksanen, 2001; Hoset, Kyrö, Oksanen, Oksanen, & Olofsson, 2014 for details on index trapping protocol). As the number of *M. agrestis* individuals from JRA was not sufficient, we supplemented the JRA samples with individuals (n=40) trapped in comparable heath, meadow and snowbed habitats at Skillefjordnes (see Ruffino et al., 2015 for site description). We also included three *M. rufocanus* individuals form this site. Altogether, WF fecal samples included 107 *L. lemmus*, 72 *Myodes rufocanus*, 103 *M. rutilus*, 93 *Microtus oeconomus* and 97 *M. agrestis* (Supplementary Table S1a).

Another set of fecal samples (n = 85) came from animals trapped in three river catchments in East Finnmark – Ifjordfjellet (70°N, 27°E), Komagdalen and Vestre Jakobselv (70-71°N, 28-31°E) (hereafter East-Finnmark or “EF”). Similarly to WF, trapping was conducted bi-annually in 15 by 15m quadrats, but samples were only collected in 2015. A total of 80 quadrats are distributed in heath and meadow habitats. The EF samples included mainly *Microtus oeconomus* (n=42) and *Myodes rufocanus* (n=38), with two *L. lemmus* and one *Microtus rutilus* (Supplementary Table S1a).

All rodents were frozen after trapping, and information of trapping date and sampling quadrant was recorded. We also recorded individual rodent body mass, sex, species identity and female reproductive status (visible pregnancy). Dataset included individuals of varying age based on the wide body mass distribution (*L. lemmus* 32-106 g, *M. rufocanus* 8-65 g, *M. rutilus* 12-36 g*, M. oeconomus* 22-90 g and *M. agrestis* 17-66 g).

Feces were carefully sampled from the intestine without damaging or breaking the intestinal tissue, avoiding contamination with blood or other compounds that could affect the NIR-spectra. From each individual from WF, we collected consecutive pellets in two Eppendorf tubes, to form two parallel samples, while of EF samples no parallel samples were taken. All pellets were dried at 40°C for minimum of 48h or until dry, and stored in Eppendorf tubes in room temperature. One parallel WF sample and all EF samples were NIR-scanned without exposure treatment (intestinal samples), while the other WF parallel sample was scanned after the exposure treatment (exposed samples).

### Exposure treatment

We subjected one set of parallel WF samples to ambient weather conditions (hereafter “exposure treatment”) in Tromsø, Northern Norway during autumn 2014 to incorporate the effects of leaching and bleaching due to UV-radiation on NIR-spectra in the calibration model. Only four species were included in the weathering treatment, as *Myodes rutilus* pellets were not suited for weathering due to a frequently liquid consistency and small size of the pellets. We placed the pellets on wooden frames of 50x50 cm, with a 1 x 1mm nylon mesh in the bottom. Nylon mesh was used as a bottom material because it does not release chemicals to the faeces and allows for water drainage and evaporation. Each frame had pellets of only one species to avoid contamination. Each frame had ca. 150 pellets. For *L. lemmus* and *Myodes rufocanus* pellets we used two frames per species, whereas *Microtus oeconomus* and *M. agrestis* had one frame each. Frames were placed on heath vegetation in September, and weather conditions during the treatment were variable, including rain, sunshine, frost and temperatures both above and below zero. During six weeks, we weekly took ca. 25 pellets for scanning from each frame.

### NIRS scanning of the fecal pellets

We scanned individual intestinal and exposed samples without further sample pre-preparation. Due to the small size, we used a custom built, Ø4mm sample holder and pressed the pellet flat with an Ø4mm metal rod in order to prevent light from penetrating through the sample. A few samples were too small to cover the whole area, but this did not affect the shape (and hence representativeness) of the spectra based on visual evaluation.

NIRS spectra were collected using a FieldSpec 3 (ASD Inc., Boulder, Colorado, USA): each spectrum was recorded as reflectance using monochromatic radiation at 1.4 nm intervals in the 350-1050nm range and 2nm intervals in the 1050-2500nm range, and interpolated to 1nm resolution. We scanned each sample three times by rotating the sample holder between scans in order to account for angular effects on light scattering. We used the mean of the three spectras in the further analysis. We used the R package *prospectr* (Stevens & Ramirez-Lopez, 2015) for spectra pre-processing, i.e. applying splice correction and normalizing the spectra. The spectral regions in the 350-399nm and 2451-2500nm ranges were removed from the dataset due to instrumental noise.

### DNA-metabarcoring of rodent feces

After NIRS scanning, we analysed a subset of intestinal WF samples (n=385) for diet species composition with DNA-metabarcoding. All wet laboratory work was performed by Center of Evolutionary Applications, University of Turku, Finland. We used a DNA metabarcoding approach similar to Soininen et al (2015). We here give a summary of the methods, and a detailed description is included in supplementary materials (p. 2: Detailed DNA-metabarcoring of rodent feces). DNA was first extracted, and thereafter amplified using universal plant primers ‘g-h’ and ‘c-h’ that target chloroplast trnL intron (Taberlet, Gielly, Pautou, & Bouvet, 1991; Taberlet et al., 2007). Final DNA libraries were constructed in the second PCR (see e.g. Vesterinen, Puisto, Blomberg, & Lilley, 2018). Resulting DNA libraries were purified and size-selected using a SPRI bead clean-up (see Vesterinen et al., 2016), concentrations were measured by Qubit (Invitrogen; www.invitrogen.com) and finally pooled in equimolar quantities. Sequencing was done on 316 v2 and 318 v2 chips with Ion PGM (Life Technologies, manual cat nr 00009816, Rev C.0).

The four sequencing runs yielded altogether 12,575,869 quality-controlled reads that could be assigned to original samples. The reads were uploaded to CSC servers (IT Center for Science, www.csc.fi) for trimming and further analysis. Bioinformatic steps were carried out following Schmidt et al. (2018) with several modifications (p. 3-4: Bioinformatics). To summarise, reads were merged, filtered for quality, cut for primers, collapsed into unique haplotypes, and finally clustered into zero-radius OTUs using softwares *usearch* (Edgar, 2010) and *cutadapt* (Martin, 2011). We were able to map 7,726,911 reads (∼93% of the trimmed reads) to our original samples. The *trnl* OTUs were initially identified to taxa using the *usearch* ‘*sintax*’ classifier with a database consisting of artic plant trnl sequences (Willerslev et al., 2014) using 70% probability threshold for taxonomic assignation. After this, read counts for each diet taxon in each sample were transformed to relative read abundances (RRA), as read abundances may actually be less misleading than presence/absence conversions (Deagle et al., 2018).

### Calibration modelling of genus and species identity

After obtaining NIR-spectra and diet data on individual samples, we built calibration models for genus and species identification and further used regression, multivariate and latent variable modelling in order to address our four hypothesis and test of concept (Fig. 1, Table 1). To build calibration models, we applied multivariate adaptive regression splines (MARS; Friedman, 1991) both directly and in a flexible discriminant analysis framework (FDA; Hastie, Tibshirani, & Buja, 1994; see Table 1). For this we used R packages *earth* (Milborrow, 2014) and *mda* (Hastie, Tibshirani, & Buja, 2015). We chose this approach as it deals well with high-dimensional input data and allows for additive effects and interactions between variables (Friedman & Roosen, 1995), and assumes non-linear responses or decision boundaries (Hastie et al., 1994). A more detailed description of the modelling and a comparison with other common chemometric approaches is provided in Supplementary material (p. 4, Calibration modelling of genus and species identity). We used R version 3.5.1 (R Development Core Team, 2018) for all statistical analyses and the *ggplot2* (Wickham, 2009) environment for plotting all figures.

We built separate calibration models for each hypothesis (H1-H4, Table 1). Models were evaluated using a modified, repeated double Monte-Carlo cross-validation (rdMCCV; cf. Xu et al., 2004; Filzmoser, Liebmann, & Varmuza, 2009) with 200 iterations. Within each iteration, we divided the modelling dataset into a *calibration set* dataset used for model fitting and variable selection and an *MCCV test set* for an assessment of model prediction accuracy, i.e. of the generalization error of each iterated calibration model (reported as model misclassification rate and species prediction accuracy; Pasquini, 2003; Filzmoser et al., 2009). The main purpose of applying rdMCCV was robust hypothesis testing, i.e. to estimate how model performance and misclassification rates vary across sets of calibration data (Filzmoser et al., 2009), and to estimate error rates for individual samples (cf. Liu, Cai, & Shao, 2008). Furthermore, we repeated the rdMCCV procedure with 95% / 5%, 90% / 10% and 80% / 20% data splits between calibration and test sets. This increased the number of total rdMCCV iterations per model to 600 (Table 1). To ensure allocation of targeted variability between calibration and test sets, the data was split (depending on the model) as fixed percentage of each species; species and exposure treatment duration; or species, exposure treatment duration and region (Table 1). We used MARS algorithms with one-way or two-way interactions of model variables, i.e. hinge functions of NIR-spectra (Table 1), depending on which interaction structure provided the lowest misclassification rate of test samples based on preliminary models with 20 iterations (Filzmoser et al., 2009).

To test hypothesis H1 (*Can we predict genus or species identity based on their fNIR-spectra?*), we built calibration models using WF intestinal and exposure sample spectra (Table 1), and constructed separate models for species and genus-level identification. Model response variable had three levels in the genus model and five levels in the species model (Table 1). In our dataset, the genus level *Lemmus* consists of only one species. In spite of this, we included the taxon in both species and genus level models, to systematically assess model predictive performances. To ensure comparability with H2 model (see below), we ran and additional rdMCCV calibration model without *Myodes rutilus*. This did not change results for the other four species, and results are not included.

To test hypothesis H2 (*Does feces exposure reduce prediction accuracy?*), we built calibration models for genus and species identity with data from the exposure treatment only. Modeling follows the description for H1, with deviations described in Table 1. We excluded *Myodes rutilus*, as the species was omitted from the exposure treatment. We then tested if individual sample prediction accuracy was explained by exposure time. For this, we used Bayesian regression models on species identity results of H1 and H2 models separately for each species and model. As response variable, we used the *individual-sample prediction accuracy* (hereafter IPA). To calculate this, we extracted the results per each individual in each model iteration, with “0” denoting a misclassification, and “1” a correct classification. We then averaged classification result across all iterated model runs. Prior to analysis, we transformed the IPA values as

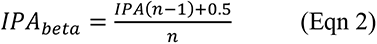

to avoid zero-one inflated data (Smithson & Verkuilen, 2006). As predictor variable we used exposure treatment time (number of weeks from 0 to 6 as a numerical fixed factor). Due to the beta-distribution (0 < y < 1) of IPA values, we fitted beta-regressions with a logit-link using package *rstanarm*, and with the function’s default weakly informative priors (Muth, Oravecz, & Gabry, 2018). We fitted the model using Markov chain Monte Carlo sampling (MCMC) with 4 chains and 2000 iterations in each chain (rstanarm default; Muth et al., 2018). We checked sampling quality of draws from the target posterior distribution for MCMC via numerical checks (R̂ < 1.1 and neff > 1200 for all model parameters) and visually with trace plots, and we inspected and confirmed good model fits by visual posterior predictive checking (Muth et al., 2018) with package *bayesplot* (Gabry & Mahr, 2019).

To test of hypothesis H3 (*Does diet confound prediction accuracy?*), we based our inference jointly on two different modelling strategies. First, we assessed the correlation between fNIR and diet RRA multivariate datasets using Mantel test with Bray-Curtis distance and Procrustes test (Jackson, 1995) using package *vegan* (Oksanen et al., 2018). While Mantel test is widely used, it may fail to account for the mean-variance relationship of the diet data (Warton, Wright, & Wang, 2012); hence, we used the Procrustes test in addition. Second, we visually explored whether highly misclassified individuals (IPA < 0.60) displayed deviant diet composition compared to correctly classified individuals of the same species. The IPA-values used here were derived from calibration models built as described for H1, but including only those intestinal samples used for DNA-metabarcoding (n=380; Table 1). To discern patterns of intra- and interspecific diet similarity, we applied multivariate latent variable modelling (model-based unconstrained ordination; Warton et al., 2015) on diet RRA data. We fitted the model using Bayesian Markov Chain Monte Carlo estimation within the HMSC modelling framework (package *Hmsc*, Blanchet, 2013; Ovaskainen et al., 2017). To estimate the model (latent) parameters, we used two MCMC chains each with 15000 iterations with a burn-in of 5000, and model default priors (see Ovaskainen et al., 2017). MCMC sampling quality of posterior draws was checked numerically and was deemed good (R̂ < 1.1 and n_eff_ > 1750). We then plotted the 2D latent variable plot of diet composition, and added the information of sample IPA-values in the plot.

To test hypothesis H4 (*Are models generalizable between regions?*), we used *Microtus oeconomus* and *Myodes rufocanus* data from both WF and EF, as these were the only numerous species in EF. First (H4.1a,b), we built separate calibration models based on data from either WF or EF only, and predicted species identity of test data including both regions (see Table 1). Thereafter (H4.2), we built calibration models with WF and EF data, and predicted MCCV test set individuals from both regions.

Finally, we demonstrate the practical application of our calibration model and the rdMCCV framework. This includes building a calibration model (Step 1), predicting new samples not seen by the calibration model (Step 2) and visualizing the calibration model predictions and misclassification risk of WF and EF samples in the FDA-model discriminant space (Step 3). Methods and results for each step are described in Box 1.

#### BOX 1

##### Practical application of the calibration model

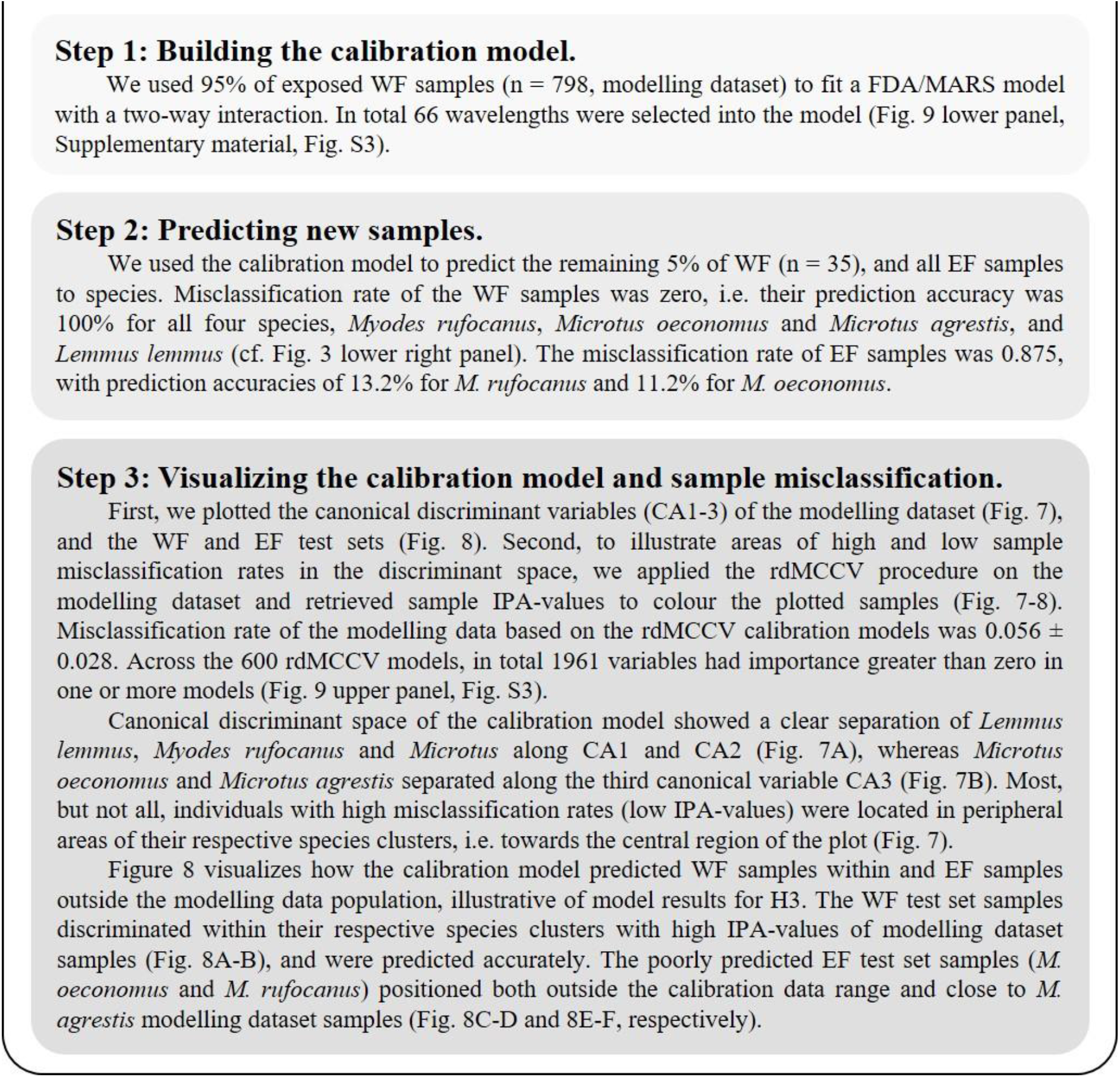

## RESULTS

### H1: Prediction accuracy of genus and species identity

Calibration models for genus identity resulted in excellent prediction accuracy and a low misclassification rate of 0.041 ± 0.019 (mean ± sd, Figure 2, upper right panel). Across all model iterations, prediction accuracy for *Myodes* was 96.2% ± 3.13%, for *Microtus* 95.1% ± 3.62% and for *Lemmus* 96.7 ± 3.47% (Fig. 2, upper left panel). For species, the prediction accuracy was more variable and with a misclassification rate of 0.129 ± 0.032 (Figure 3, upper right panel). Prediction accuracies were high for *Myodes rufocanus* (95.4% ± 4.02%), *Myodes rutilus* (91.6% ± 9.20%) as well as *Lemmus lemmus* (96.1% ± 3.18%), but much poorer for *Microtus oeconomus* (68.8% ± 10.7%) and *Microtus agrestis* (75.1% ± 9.85%) (Fig. 3 upper left panel).

**Figure 2.**
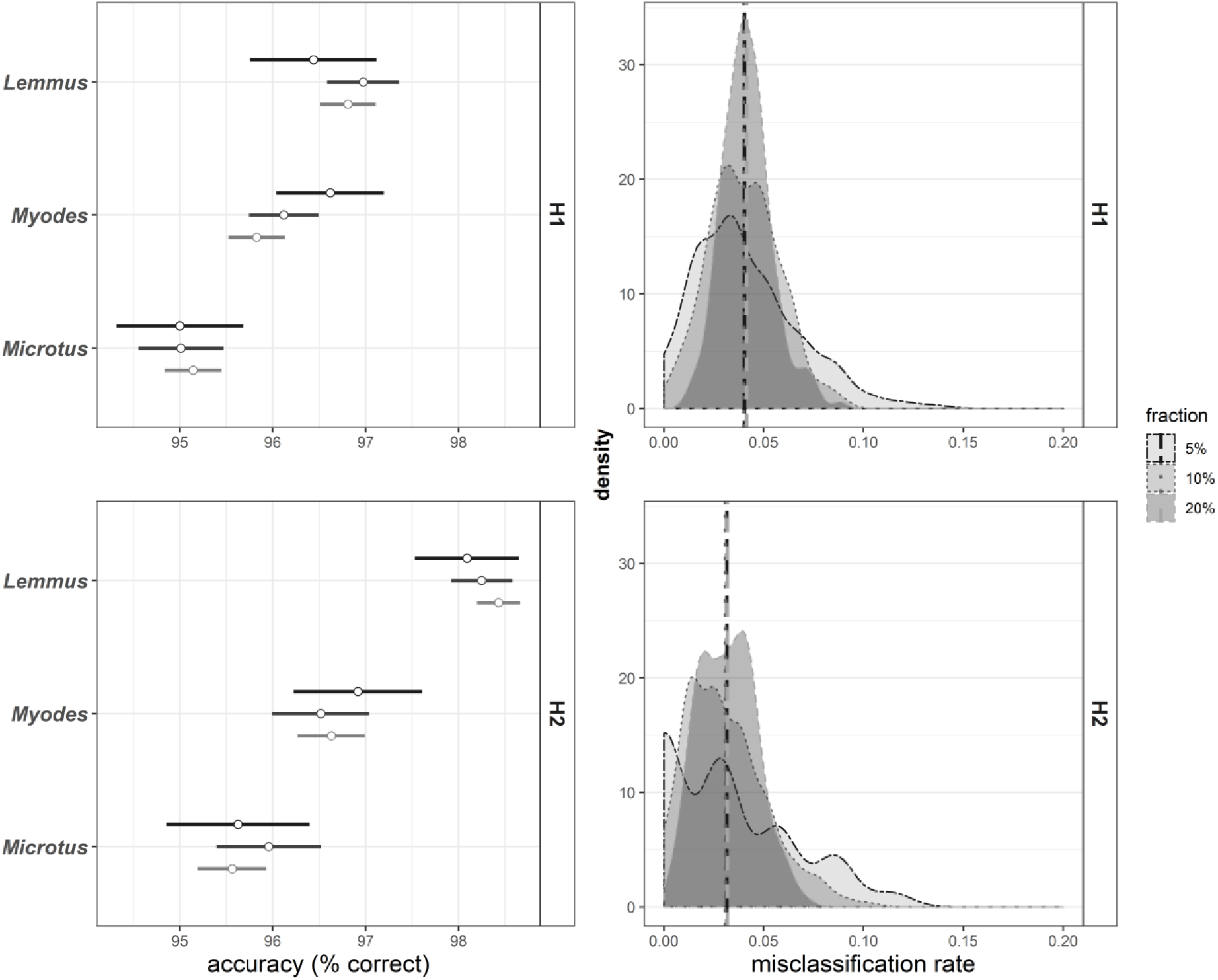
Predictive ability of genus-specific fNIRS calibrations for small rodents from Finnmark, Norway. Left panels: prediction accuracy of each genus (mean ± sd) separately. Right panels: density plots of model misclassification rates. Upper row shows results from calibration models used for H1 (i.e. including all data from West Finnmark), lower row for H2 (i.e. including only samples used to test exposure effect). Results are divided by data splits (fractions) with 200 iterations each, where 5%, 10% and 20% of data was randomly assigned as validation data (using rdMCCV framework).

**Figure 3.**
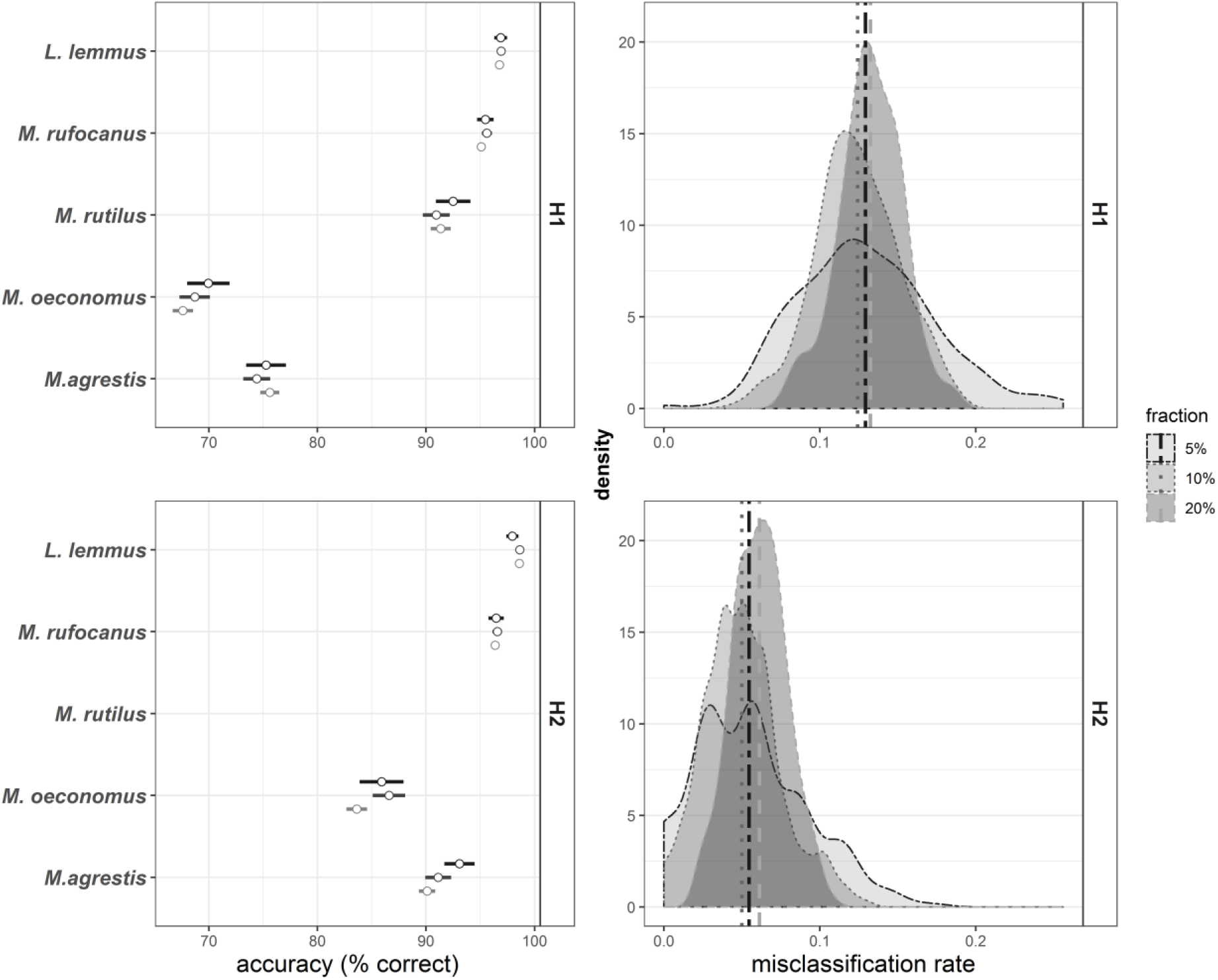
H1 (upper row) and H2 (lower row) species FDA-model results. Results are divided by data splits (fractions) with 200 iterations each, where 5%, 10% and 20% of data was randomly assigned to the MCCV test set. Left panels: prediction accuracy of each species (mean ± sd) separately. *M.rutilus* samples were not included in exposure treatment and H2 model. Upper panels: density plots of model misclassification rates.

### H2: effect of feces exposure

Calibration models based on only exposed samples performed better than calibration models including intestinal samples, especially in predicting species identity (Fig. 3). Effect of feces exposure to ambient weather for 1-6 weeks was visible on fNIR-spectra, with a clear decline in reflectance at 700-1400nm, and increase in reflectance between ca. 1500-2500nm (Supplementary material, Fig. S1). The rate of change in fNIR-spectra seemed to decline with exposure duration for all species; however the *Microtus oeconomus* and *M. agrestis* spectra changed strongly again after week 5 (Supplementary material, Fig. S1). Model misclassification rate for genus identity was 0.031 ± 0.02 (Fig. 2, lower right panel), while mean prediction accuracy for *Myodes* was 96.7% ± 3.89%, for *Microtus* 95.7% ± 4.24% and for *Lemmus* 98.3 ± 2.86% (Fig. 2, lower left panel). Model misclassification rates for species calibrations were low, with 0.055 ± 0.028, and mean prediction accuracies reached 96.5 ± 3.76% for *Myodes rufocanus*, 98.4% ± 2.79% for *Lemmus lemmus* and was as high as 85.4% ± 11.2% for *Microtus oeconomus* and 91.4% ± 8.25% for *Microtus agrestis* (Fig. 3, lower row).

Posterior distributions of exposure week effects on IPA-values indicated a positive effect of exposure on *Microtus oeconomus* and *M. agrestis* prediction accuracy when both intestinal and exposed samples were included in calibration (H1 model). However, exposure did not affect IPA-values when only exposed samples we used in calibration (H2 model). Exposure treatment did not affect individual prediction accuracies of other species with 95% confidence limits (Fig. 4, Supplementary Table S3).

**Figure 4.**
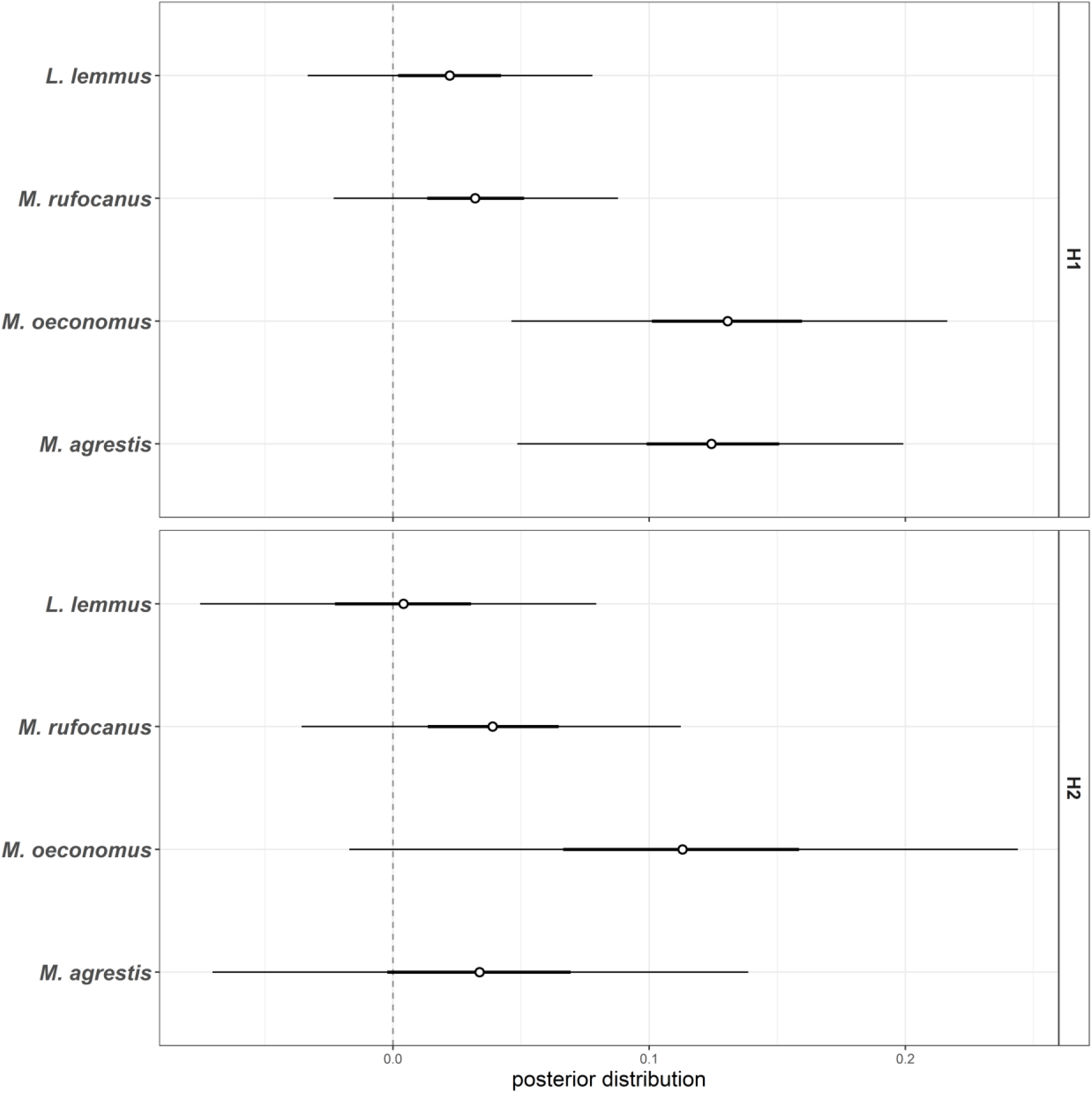
Beta-regression posterior distributions (mean ± 95% CI) of exposure duration (in weeks) effect on individual prediction accuracy (IPA) of four species included in exposure treatment. Upper panel shows model results where IPA-values are based on H1 calibration, whereas in lower panel IPA-values are based on H2 calibration.

### H3: Does diet confound prediction accuracy?

The fNIR-spectra was only weakly correlated to diet, as indicated by both Mantel test (r = 0.153, p < 0.001) and Procrustes test (m12-squared = 0.932, correlation = 0.261, p < 0.001). Latent variable modelling indicated clear overlap between most species’ diets. However, *Myodes rufocanus* and *M. rutilus* were clearly differentiated (Fig. 5). Most of the individuals with low IPA values (i.e. individuals that were often classified to wrong species) were located among the dense diet cluster of their respective species, while many peripheral individuals in terms of diet had high IPA scores (Fig. 5).

**Figure 5.**
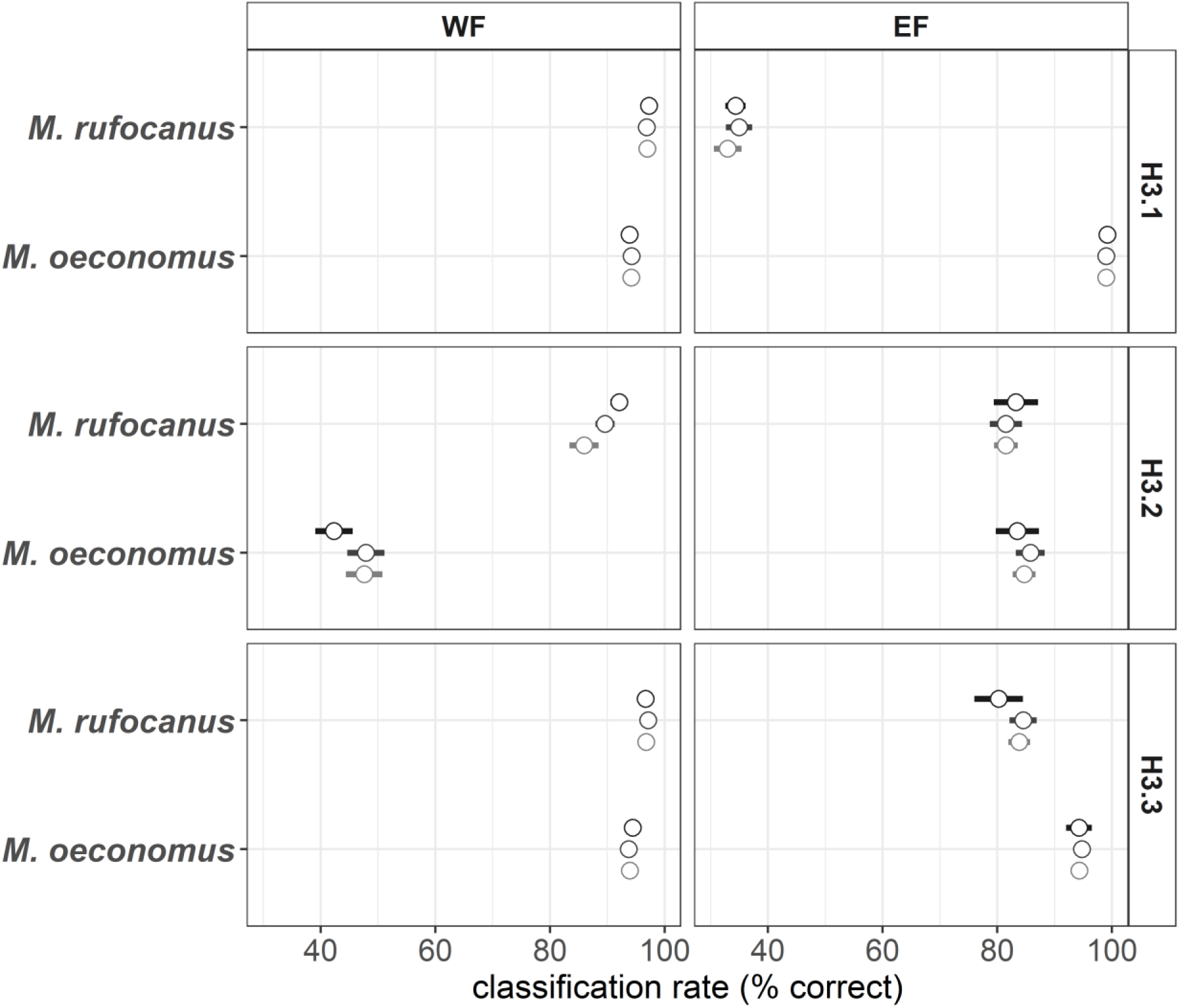
H3 model prediction accuracy (mean ± sd) for WF and EF separately. Upper row: H3.1 calibration is based on WF data only. Middle row: H3.2 calibration is based on EF data only. Lower row: H3.3, calibration is based on jointly WF and EF data.

### H4: regional vs. cross-regional calibrations

Calibration models built with data from one region predicted samples from the same region well, but samples from the other region poorly (Fig 6; cf. Box 1 and Fig. 8). WF calibration (H4.1a) predicted WF samples with a misclassification rate of 0.041 ± 0.027, while misclassification rate for EF samples was as high as 0.349 ± 0.071 (Supplementary material, Fig. S2). Mean prediction accuracy of *Myodes rufocanus* was 96.8% ± 3.14% in WF samples, but only 27.1% ± 15.6 in EF samples, while prediction accuracy of *Microtus oeconomus* samples from WF was high (94.2% ± 5.30%), but not as high as in EF samples (99.4% ± 2.70%) (Fig. 6 upper row).

**Figure 6.**
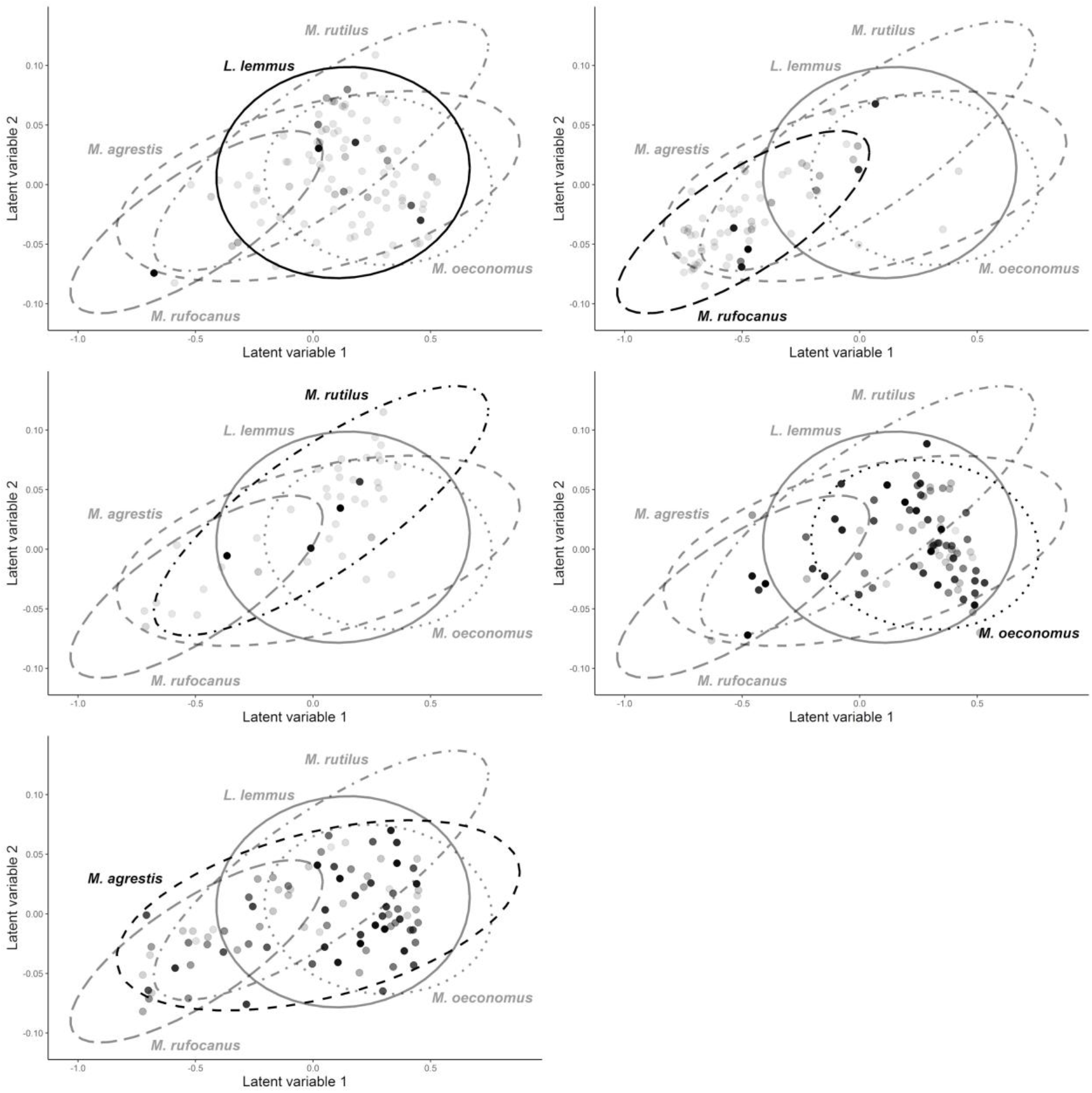
Comparing variation in diet composition with individual sample misclassification rate. Each individual (sample) is plotted as a separate dot, and all five species are plotted in their separate subplots. Spreads of species-specific diets are shown by separate ellipses. Point color darkness indicates lower IPA-value, i.e. higher sample misclassification rate based on rdMCCV calibration models on plotted samples. Point locations are latent variables of relative read abundances. Latent variable modeling is equivalent to unconstrained ordination, i.e. proximity of samples in two-dimensional space indicates diet similarity, and distance indicates dissimilarity.

Similar region-specific pattern emerged with calibration models built on EF data (H4.1b). Misclassification rate was lower for EF (0.171 ± 0.140) than for WF samples (0.265 ± 0.064, Fig. S2). Prediction accuracy for *M. oeconomus* form WF was especially poor (47.9% ± 20.2%), while *M. rufocanus* reached higher prediction accuracy (89.7% ± 12.8%) (Fig. 6 middle row). In contrast, both *M. oeconomus* (84.9% ± 19.6%) and *M. rufocanus* (80.8% ± 21.4%) from EF test data reached decent prediction accuracies (Fig. 6 middle row).

Calibration model including both WF and EF samples (H4.2), predicted test samples nearly as well or better than either of the regional models alone (Fig. 6). Misclassification rate for WF samples was 0.041 ± 0.027, and for EF samples, 0.104 ± 0.108 (Fig. S2). Mean prediction accuracy for *M. rufocanus* was 96.7% ± 3.25% for WF and 85.3% ± 18.5% for EF samples (Fig. 6 lower row). Models predicted *M. oeconomus* at 94.6% ± 5.02% and 93.9% ± 13.2% in WF and EF samples, respectively (Fig. 6 lower row).

## DISCUSSION

Our results demonstrate fNIRS as an accurate method for predicting vole and lemming genus and species identity based on single fecal pellets (H1). While the calibration models predicted genus identity extremely well (at > 95% accuracy), prediction accuracy of species identity varied more but was still good-to-excellent at 85-98%. Importantly, exposure of samples to ambient weather improved the calibration model performance, contradicting the hypothesised negative effect of collecting samples from the field (H2). Surprisingly, we did not find support for diet being a major determinant of species prediction accuracy (H3). Furthermore, while our results indicate that calibration models based on samples from one region may not readily be applicable to samples from another region, calibration models including samples from two regions provide good prediction accuracy for both regions, corroborating our fourth hypothesis (H4). These findings are in line with previous results of reaching good prediction accuracies when calibration datasets are expanded in space or time (Douglas R. Tolleson et al., 2005; Murguzur et al., 2019). Our results lend strong support for using fNIRS on samples exposed to ambient conditions, i.e. samples collected from the field, and for further development of fNIRS for large-scale rodent population monitoring, including cross-regional or global calibrations.

### Diet and misclassification

Dried feces consist of ingested diet material, gut microbes and sloughed tissue, gastric secretions and metabolized hormones, of which all may carry species-specific chemical or structural imprints. Of these, we explicitly addressed the link between diet composition and misclassification. We did not find strong evidence of diet composition explaining individual sample misclassification. This result is surprising as diet quality measures such as fecal nitrogen, neutral and acid detergent fiber (NDF, ADF), crude fiber, lignin, ether extracts and ash are well predictable from fNIR spectra (Steyaert et al., 2012; Gil-Jiménez, Villamuelas, Serrano, Delibes, & Fernández, 2015). If diet quality and composition are only loosely associated, the link between fNIR spectra, species identity and generalist diet composition may not be easily detected. Even so, it is likely that group-specific variation in diet composition affecting any diet quality variable would still affect misclassification rates (e.g. Tolleson et al., 2005). Further research is needed to determine if and which diet composition or quality variables co-vary with species identity to affect fNIRS rodent species classification.

Furthermore, the large sample size from WF may have incorporated sufficiently high levels of intraspecific dietary variation, to render the effect of diet on misclassification rate small, compared to other fecal constituents. With lower sample sizes, systematic variation in diet e.g. between years could confound species identification as found by Tolleson *et al*. (2005). Similarly, systematic variation in species diet between regions could explain high rates of misclassifications between WF and EF. Species-specific diets between these regions may differ given different rodent species guild composition, habitat composition and forage availability, which could translate to variation in dietary quality. With WF calibrations, both *M. rufocanus* and *M. oeconomus* from EF were frequently misclassified, as they appeared to cluster together with *M. agrestis* – a species absent from Eastern Finnmark. It is possible that in the absence of *M. agrestis*, which competes with both *M. oeconomus* and *M. rufocanus* at WF, these latter two species’ dietary niches in EF resemble that of *M. agrestis* at WF.

### Juxtaposing misclassification with phylogenetic history

In general, classification model performance emerges likely as a complex product of individual traits linked with phylogenetic history. Evolutionary constraints may manifest in fNIR-spectra through morphological, physiological or microbiological differences in the digestive system, along with ecological niche factors (Blomberg, Garland, & Ives, 2003). Dental and gut morphologies are examples of phylogenetically variable traits that may affect fNIRS species identification through differences in fecal particle size (Sheine & Kay, 1977; Clauss et al., 2015) and fecal nitrogen levels or fibre digestibility (Lovegrove, 2010; Clauss et al., 2015), respectively (Foley et al., 1998; Tolleson et al., 2005 and references therein; Steyaert et al., 2012).

Indeed, divergence of *Lemmus*, *Myodes* and *Microtus* at tribal level (Buzan, Krystufek, Hänfling, & Hutchinson, 2008) is congruent with good performance of calibration models at genus level. Similarly, *Myodes rufocanus* and *M. rutilus* which were well separated in classification models, show marked phylogenetic differentiation within the genus (Cook, Runck, & Conroy, 2004; Buzan et al., 2008; Kohli et al., 2014). While the two *Myodes* species’ ecology can be similar in e.g. sub-arctic birch forests (cf. Ehrich, Yoccoz, & Ims, 2009), clear behavioural niche differences may exist (Nations & Olson, 2015) and for instance their habitat use at JRA differs markedly (M. Tuomi et al unpublished data). In contrast, higher misclassification rates and similarities in *Microtus* fNIR-spectra link with the relatively recent and rapid radiation history of the genus (Barbosa, Paupério, Pavlova, Alves, & Searle, 2018). While *M. oeconomus* is ancestral to *M. agrestis*, there is ongoing intraspecific divergence within both species (Barbosa et al., 2018), as well as strong interspecific competition (e.g. Hoset & Steen, 2007). Divergence patterns in dental structure converge roughly with these phylogenetic patterns, with *Microtus oeconomus* and *M. agrestis* displaying the smallest differences (Herrmann, 2002).

### Identification of species and pellet exposure

Surprisingly, misclassification rates of our samples decreased after exposure to ambient weather. This was mainly due to higher prediction accuracy within the genus *Microtus*, when we excluded intestinal samples from modelling. It appears that species-specific signals in fNIR-spectra of the closely related *M. oeconomus* and *M. agrestis* became more apparent with exposure. While we found no evidence of reduced prediction accuracy with increasing exposure time, rates of visible changes in spectra varied between weeks and species. Yet, it is likely that exposure times longer than 6 weeks will eventually lead to loss of species-specific signals as the fecal pellets decompose. Further research is needed to determine the maximum or optimal timeframe for successful classification of field-sampled rodent feces.

Juxtaposing calibrations built on intestinal and exposed samples allows for speculation as to which fecal constituents may have contributed to species identification, based on constituents’ known susceptibility to exposure. Specifically, species-specific signals in fNIR-spectra seem not to associate with volatile or rapidly leaching substances. For instance, stability of constituents linked with dietary quality in fecal samples varies strongly under exposure (Jenks et al., 1990; Leite & Stuth, 1994; Kamler et al., 2003; Steyaert et al., 2012). The fecal metabolome likely displays species-specific variation (Zierer et al., 2018) detectable by fNIRS (Saric et al., 2008; Santos et al., 2014), yet fecal steroid metabolite concentrations of various large mammal species decline during days or remain stable for only up to a week (Abáigar, Domené, & Palomares, 2010; Mesa-Cruz, Brown, & Kelly, 2014; Parnell et al., 2015). Thus, steroid metabolites may be too short-lived to have accounted for species identification in our study. In contrast, differences in microbial flora could translate to some exposure-resistant differences in rodent fNIR-spectra. For instance, diaminopimelic acid (DAPA), a marker of gut microbe derived N in feces (Karr-Lilienthal et al., 2004) has good NIRS calibrations (Atanassova, Todorov, Djouvinov, Tsenkova, & Toyoda, 1998), and has been found to retain stable concentrations in exposed deer feces (Kamler et al., 2003). In summary, rodent species identification is likely to rest on a multitude of fecal constituents, many of which link with phylogenetic distance (Ley et al., 2008; Zierer et al., 2018), and have varying resistance to exposure and decay. We note that quoted studies on exposure effects on fecal dietary quality indicators and on fecal metabolome cover only up to a few weeks or days, respectively, providing only a preliminary basis for interpretation of constituents affecting fNIRS calibrations.

### Towards application of fNIRS calibrations for rodent monitoring

Practical application of fNIRS for monitoring and census purposes rests on good understanding of model behaviour and limitations of calibration models. Notably, we found that the selected wavelengths varied considerably between model iterations (see Box 1), thus providing little support for any definite set of wavelengths being superior for species identification. Instead, our modelling framework provides a visual tool to assess misclassification risk of new samples derived from the calibration data population (Figs. 7-8). Samples located well within species clusters in the discriminant space are likely to be predicted correctly, while samples falling between the clusters have a considerably higher risk of misclassification. Such samples should optimally be used to improve model decision boundaries by including them in the model calibration. To control for inter-annual variation or drift in e.g. diet in long-term monitoring, a small subset of new samples should, at regular intervals if possible, undergo an external verification of species identity, and thus compliment the calibration dataset.

**Figure 7.**
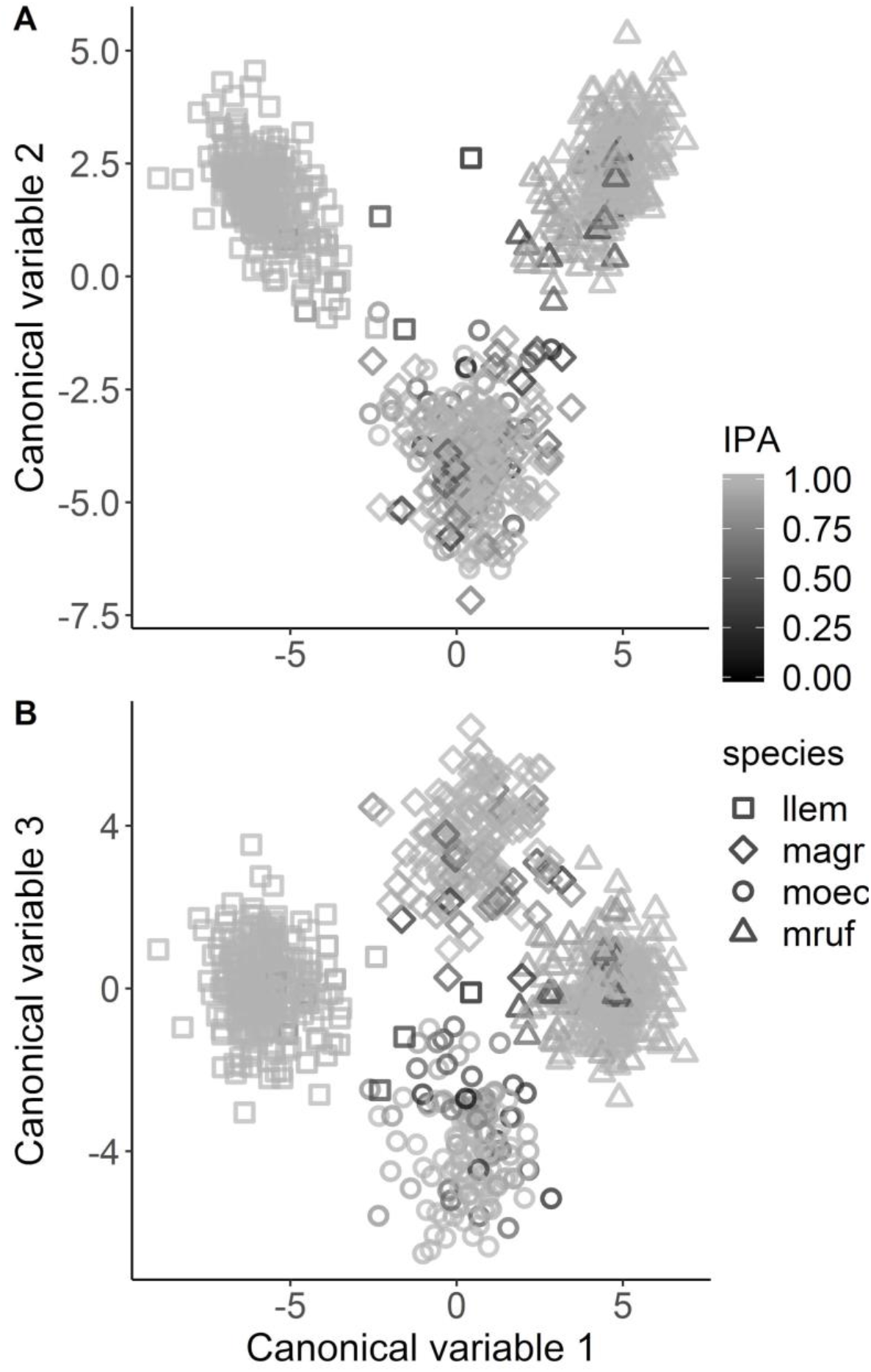
Canonical discriminant plot of the calibration model. Eeach point indicates modeling dataset sample position in the FDA model’s 3D canonical discriminant space. Point shape indicates species identity, and point shade of grey indicates the individual sample IPA-value so that hue darkness denotes increasing misclassification rate. llem = *Lemmus lemmus*, mruf = *Myodes rufocanus*, moec = *Microtus oeconomus*, magr = *Microtus agrestis*.

**Figure 8.**
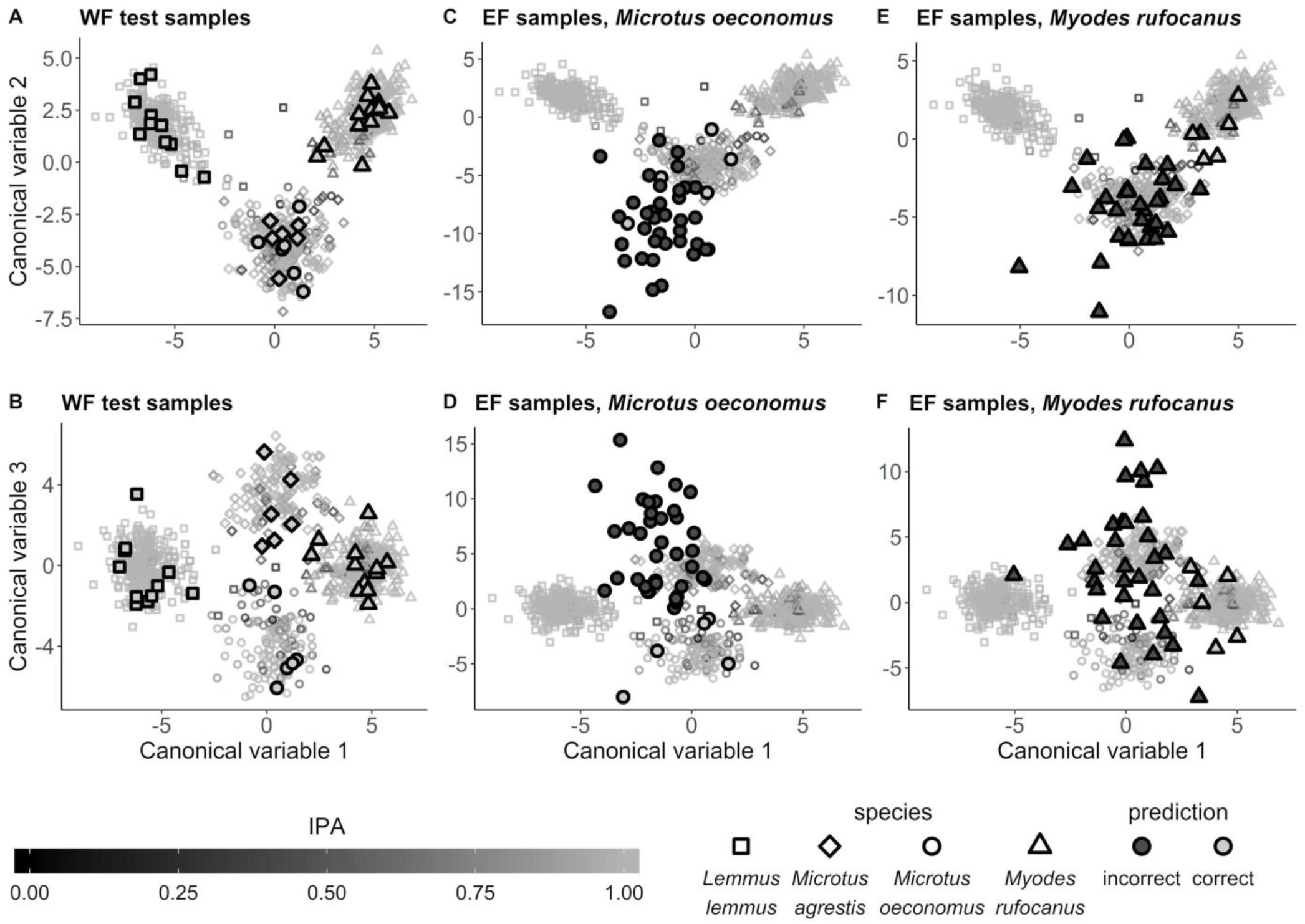
Canonical discriminant plot of the calibration model, with predicted locations of WF and EF test samples plotted on top of the modeling dataset points (cf. Fig. 7). Point shape indicates species identity. WF and EF samples are plotted with large points with black borders, and with fill colour indicating correct or incorrect calibration model prediction. Modeling data are plotted in all panels as small transparent grey points as in Fig. 7.

**Figure 9.**
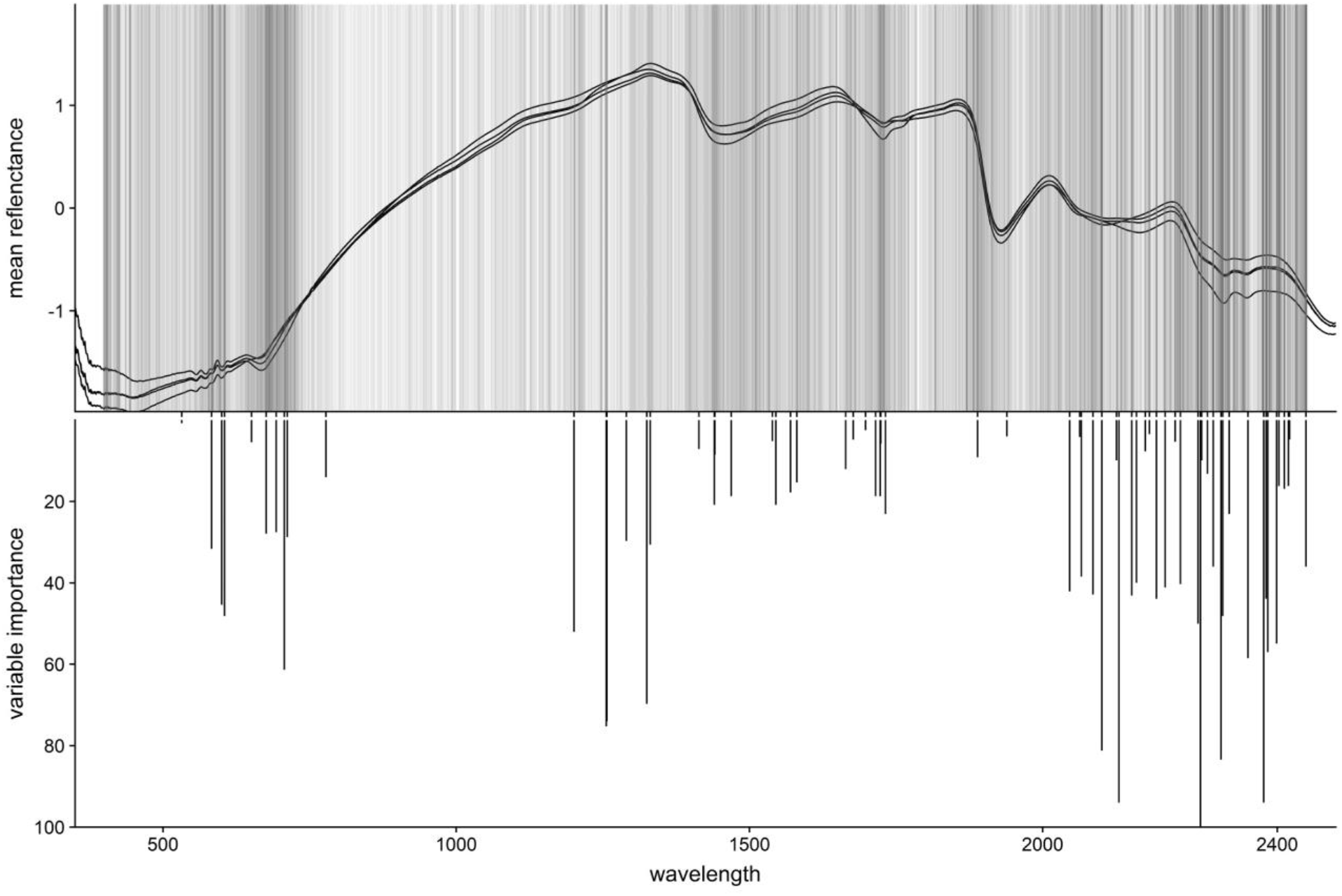
Comparison of the calibration model’s variable selection and variable importance (VI) with the variable selection across all rdMCCV models. Wavelenght is plotted on the x-axis. **Upper panel:** Wavelenghts with VI > 0 across all rdMCCV iterations. Increasing hue darkness indicates higher frequency of VI > 0 for that wavelength across all 600 rdMCCV iterations (a grey, transparent vertical line is plotted each time a wavelength receives a VI score > 0). Plot includes mean spectra of all four species in the calibration model, to visualize differences in spectra with important wavelength distribution. **Lower panel:** Wavelenghts with VI > 0 in the calibration model, with VI score on the reversed y-axis.

Our results corroborate previous findings of high or increased misclassification rates of samples from other regions or years (Tolleson et al., 2005; Murguzur et al., 2019), and of breaking past limitations of closed sample populations (Murguzur et al., 2019). New samples from external populations may exhibit considerably different variation in constituents underlying the species-specific signal compared to calibration samples, making their predictive modelling highly risky. However, we show that increasing the spatial extent of calibration data will produce robust models across sample populations. This is hardly surprising, if the species-specific signal is co-determined by a complex host of constituents, some of which are linked to community-specific interactions (e.g. diet, body condition), while others would be more unequivocally associated with phylogenetic constraints. Therefore, increasing the variability in calibration data should mask constituents linked with community-specific interactions, and lead to detection of wavelength combinations predictive of species identity across populations. While diet, habitat use and gut morphology of microtine rodents show high levels of plasticity (Lovegrove, 2010; Soininen et al., 2013), a growing body of fNIRS research (Douglas R. Tolleson, 2010; Vance, Tolleson, Kinoshita, Rodriguez, & Foley, 2016; Aw & Ballard, 2019) suggests that robust species-specific markers are also likely to exist.

In conclusion, fNIRS can facilitate rodent population censuses at larger spatial extent and smaller grains than before deemed feasible, if combined with pellet-count based abundance indices (Karels et al., 2004; Engeman & Whisson, 2006; Jareño et al., 2014). The wide array of data (e.g. diet, disease, stress) discernible through fNIRS suggests that developing monitoring schemes based on pellet counts and fNIRS could meet the need for ecosystem- and interaction-based approaches to monitoring (Ehrich et al., 2019). Further steps in this direction involve increasing the spatial scope of calibrations, extending the calibration exposure times to meet the needs of field sampling intervals, as well as continuous development of the calibration model algorithm and outlier detection. Key advantage of fNIRS is the ability to use existing spectral data to develop calibrations for any number of qualitative and quantitative constituents (Foley et al., 1998; Vance et al., 2016), opening possibilities to link process and pattern across different levels of organization. Thus, given the emergence of active global researcher networks (e.g. Barrio et al., 2016) and spectral libraries (e.g. Shepherd & Walsh, 2002), we propose initiatives towards development of circumpolar, -boreal or global rodent fNIRS calibrations.

## Acknowledgements

NIR and diet metabarcoding analyses were funded by FRAM Center (grant to KAB). British Ecological Society (BES) provided additional funding for the diet analysis (grant to KSH). Small rodents were obtained after data collection for long-term time series, and the animals utilized in this study were part of work within NCoE Tundra at WF and the Climate-Ecological Observatory for Arctic Tundra (COAT) at EF. The work was financed by Turku University Foundation, the FRAM Center and the Research Council of Norway through the Climate-Ecological Observatory for Arctic Tundra (COAT). We thank Riikka Rinnan for her comments on the manuscript version. We especially thank Lise Ruffino and all field personnel involved in collecting small rodents for this study, as well as Joatka fjellstue for lodging at JRA.

## Data accessibility statement

Small rodent faeces spectral data, dietary data with masked species identity and R scripts for rdMCCV framework model will be available at UiT Open Research Data (https://opendata.uit.no) should the manuscript be accepted.

## Author contributions

MT, FJAM and KAB conceived the idea and the general design; MT, KSH, FJAM, SK and TAU contributed substantially to sample collection and processing; FJAM led NIR-spectroscopy of the fecal pellets with support from MT and SK; MT and FJAM modeled the NIRS spectra; EV and ES designed and executed diet DNA-metabarcoding and bioinformatics. KAB was responsible of project administration and supervised the work of MT. KAB, KSH and MT acquired funding. MT wrote the first draft. All authors contributed substantially to the manuscript draft during multiple commenting rounds, and gave a final approval to the submitted version.

## REFERENCES

1. Abáigar, T., Domené, M. A., & Palomares, F. (2010). Effects of fecal age and seasonality on steroid hormone concentration as a reproductive parameter in field studies. European Journal of Wildlife Research, 56(5), 781–787. doi:10.1007/s10344-010-0375-z

2. Atanassova, S., Todorov, N., Djouvinov, D., Tsenkova, R., & Toyoda, K. (1998). The Possibility of near Infrared Spectroscopy for Evaluation of Microbial Nitrogen Content of in Sacco Feed Residues and Duodenal Digesta of Sheep. Journal of Near Infrared Spectroscopy, 6(1), 167–174. doi:10.1255/jnirs.133

3. Aw, W. C., & Ballard, J. W. O. (2019). Near-infrared spectroscopy for metabolite quantification and species identification. Ecology and Evolution, 9(3), 1336–1343. doi:10.1002/ece3.4847

4. Barbosa, S., Paupério, J., Pavlova, S. V., Alves, P. C., & Searle, J. B. (2018). The Microtus voles: Resolving the phylogeny of one of the most speciose mammalian genera using genomics. Molecular Phylogenetics and Evolution, 125, 85–92. doi:10.1016/j.ympev.2018.03.017

5. Barrio, I. C., Hik, D. S., Jónsdóttir, I. S., Bueno, C. G., Mörsdorf, M. A., & Ravolainen, V. T. (2016). Herbivory Network: An international, collaborative effort to study herbivory in Arctic and alpine ecosystems. Polar Science, 10(3), 297–302. doi:10.1016/j.polar.2016.03.001

6. Blanchet, F. G. (2013). HMSC: Hierarchical modelling of ecological communities. Retrieved from https://researchportal.helsinki.fi/en/publications/hmsc-hierarchical-modelling-of-ecological-communities

7. Blomberg, S. P., Garland, T., & Ives, A. R. (2003, April 1). Testing for phylogenetic signal in comparative data: Behavioral traits are more labile. doi:10.1111/j.0014-3820.2003.tb00285.x

8. Buzan, E. V., Krystufek, B., Hänfling, B., & Hutchinson, W. F. (2008). Mitochondrial phylogeny of Arvicolinae using comprehensive taxonomic sampling yields new insights. Biological Journal of the Linnean Society, 94(4), 825–835. doi:10.1111/j.1095-8312.2008.01024.x

9. Campbell, D., Swanson, G. M., & Sales, J. (2004). Methodological Insights: Comparing the precision and cost-effectiveness of faecal pellet group count methods: Faecal pellet group count methods. Journal of Applied Ecology, 41(6), 1185–1196. doi:10.1111/j.0021-8901.2004.00964.x

10. Clauss, M., Steuer, P., Erlinghagen-Lückerath, K., Kaandorp, J., Fritz, J., Südekum, K.-H., & Hummel, J. (2015). Faecal particle size: Digestive physiology meets herbivore diversity. Comparative Biochemistry and Physiology Part A: Molecular & Integrative Physiology, 179, 182–191. doi:10.1016/j.cbpa.2014.10.006

11. Cook, J. A., Runck, A. M., & Conroy, C. J. (2004). Historical biogeography at the crossroads of the northern continents: molecular phylogenetics of red-backed voles (Rodentia: Arvicolinae). Molecular Phylogenetics and Evolution, 30(3), 767–777. doi:10.1016/S1055-7903(03)00248-3

12. Deagle, B. E., Thomas, A. C., McInnes, J. C., Clarke, L. J., Vesterinen, E. J., Clare, E. L., … Eveson, J. P. (2018). Counting with DNA in metabarcoding studies: How should we convert sequence reads to dietary data? Molecular Ecology, 0(0). doi:10.1111/mec.14734

13. Dickman, C. R. (1999). Rodent-ecosystem relationships: a review. Ecologically-Based Management of Rodent Pests. ACIAR Monograph, (59), 113–133.

14. Edgar, R. C. (2010). Search and clustering orders of magnitude faster than BLAST. Bioinformatics, 26(19), 2460–2461. doi:10.1093/bioinformatics/btq461

15. Ehrich, D., Schmidt, N. M., Gauthier, G., Alisauskas, R., Angerbjörn, A., Clark, K., … Solovyeva, D. V. (2019). Documenting lemming population change in the Arctic: Can we detect trends? Ambio. doi:10.1007/s13280-019-01198-7

16. Ehrich, D., Yoccoz, N. G., & Ims, R. A. (2009). Multi-annual density fluctuations and habitat size enhance genetic variability in two northern voles. Oikos, 118(10), 1441–1452. doi:10.1111/j.1600-0706.2009.17532.x

17. Ekerholm, P., Oksanen, L., & Oksanen, T. (2001). Long-term dynamics of voles and lemmings at the timberline and above the willow limit as a test of hypotheses on trophic interactions. Ecography, 24(5), 555–568. doi:10.1111/j.1600-0587.2001.tb00490.x

18. Engeman, R., & Whisson, D. (2006). Using a general indexing paradigm to monitor rodent populations. International Biodeterioration & Biodegradation, 58(1), 2–8. doi:10.1016/j.ibiod.2006.03.004

19. Fauteux, D., Gauthier, G., Mazerolle, M. J., Coallier, N., Bêty, J., & Berteaux, D. (2018). Evaluation of invasive and non-invasive methods to monitor rodent abundance in the Arctic. Ecosphere, 9(2), e02124. doi:10.1002/ecs2.2124

20. Filzmoser, P., Liebmann, B., & Varmuza, K. (2009). Repeated double cross validation. Journal of Chemometrics, 23(4), 160–171. doi:10.1002/cem.1225

21. Foley, W. J., McIlwee, A., Lawler, I., Aragones, L., Woolnough, A. P., & Berding, N. (1998). Ecological applications of near infrared reflectance spectroscopy–a tool for rapid, cost-effective prediction of the composition of plant and animal tissues and aspects of animal performance. Oecologia, 116(3), 293–305.

22. Friedman, J. H. (1991). Multivariate Adaptive Regression Splines. The Annals of Statistics, 19(1), 1–67. doi:10.1214/aos/1176347963

23. Friedman, J. H., & Roosen, C. B. (1995). An introduction to multivariate adaptive regression splines. Statistical Methods in Medical Research, 4(3), 197–217.

24. Gabry, J., & Mahr, T. (2019). bayesplot: Plotting for Bayesian Models. Retrieved from mc-stan.org/bayesplot

25. Galan, M., Pagès, M., & Cosson, J.-F. (2012). Next-Generation Sequencing for Rodent Barcoding: Species Identification from Fresh, Degraded and Environmental Samples. PLOS ONE, 7(11), e48374. doi:10.1371/journal.pone.0048374

26. Gil-Jiménez, E., Villamuelas, M., Serrano, E., Delibes, M., & Fernández, N. (2015). Fecal Nitrogen Concentration as a Nutritional Quality Indicator for European Rabbit Ecological Studies. PLoS ONE, 10(4), e0125190. doi:10.1371/journal.pone.0125190

27. Green, M. L., Ting, T.-F., Manjerovic, M. B., & Mateus-Pinilla, N. (2013). Noninvasive alternatives for DNA collection from threatened rodents, 2013. doi:10.4236/ns.2013.55A003

28. Hastie, T., Tibshirani, R., & Buja, A. (1994). Flexible Discriminant Analysis by Optimal Scoring. Journal of the American Statistical Association, 89(428), 1255–1270. doi:10.1080/01621459.1994.10476866

29. Hastie, T., Tibshirani, R., & Buja, A. (2015). Package ‘mda’: Flexible discriminant analysis by optimal scoring. CRAN. org. Retrieved from http://www.tandfonline.com/doi/abs/10.1080/01621459.1994.10476866

30. Heisler, L. M., Somers, C. M., & Poulin, R. G. (2016). Owl pellets: a more effective alternative to conventional trapping for broad-scale studies of small mammal communities. Methods in Ecology and Evolution, 7(1), 96–103. doi:10.1111/2041-210X.12454

31. Herrmann, N. (2002). Food-specialization and structural parameters of dental patterns of Arvicolinae (Rodentia, Mammalia). Senckenbergiana Lethaea, 82(1), 153–165. doi:10.1007/BF03043781

32. Hoset, K. S., Kyrö, K., Oksanen, T., Oksanen, L., & Olofsson, J. (2014). Spatial variation in vegetation damage relative to primary productivity, small rodent abundance and predation. Ecography. Retrieved from http://agris.fao.org/agris-search/search.do?recordID=US201400139754

33. Hoset, K. S., & Steen, H. (2007). Relaxed competition during winter may explain the coexistence of two sympatric Microtus species. Annales Zoologici Fennici, 44, 10.

34. Ims, R. A., & Fuglei, E. V. A. (2005). Trophic interaction cycles in tundra ecosystems and the impact of climate change. Bioscience, 55(4), 311–322.

35. Jackson, D. A. (1995). PROTEST: A PROcrustean Randomization TEST of community environment concordance. Écoscience, 2(3), 297–303. doi:10.1080/11956860.1995.11682297

36. Jareño, D., Viñuela, J., Luque-Larena, J. J., Arroyo, L., Arroyo, B., & Mougeot, F. (2014). A comparison of methods for estimating common vole (Microtus arvalis) abundance in agricultural habitats. Ecological Indicators, 36, 111–119. doi:10.1016/j.ecolind.2013.07.019

37. Jenks, J. A., Soper, R. B., Lochmiller, R. L., & Leslie, D. M. (1990). Effect of Exposure on Nitrogen and Fiber Characteristics of White-Tailed Deer Feces. The Journal of Wildlife Management, 54(3), 389–391. doi:10.2307/3809644

38. Kamler, J., Homolka, M., & Kráčmar, S. (2003). Nitrogen characteristics of ungulates faeces: effect of time of exposure and storage. Folia Zoologica, 52(1), 31–35.

39. Karels, T. J., Koppel, L., & Hik, D. S. (2004). Fecal Pellet Counts as a Technique for Monitoring an Alpine-Dwelling Social Rodent, the Hoary Marmot (Marmota caligata). Arctic, Antarctic, and Alpine Research, 36(4), 490–494. doi:10.1657/1523-0430(2004)036[0490:FPCAAT]2.0.CO;2

40. Karr-Lilienthal, L. K., Grieshop, C. M., Spears, J. K., Patil, A. R., Czarnecki-Maulden, G. L., Merchen, N. R., & Fahey, G. C. (2004). Estimation of the proportion of bacterial nitrogen in canine feces using diaminopimelic acid as an internal bacterial marker. Journal of Animal Science, 82(6), 1707–1712. doi:10.2527/2004.8261707x

41. Kerr, J. T., & Ostrovsky, M. (2003). From space to species: ecological applications for remote sensing. Trends in Ecology & Evolution, 18(6), 299–305. doi:10.1016/S0169-5347(03)00071-5

42. Kohli, B. A., Speer, K. A., Kilpatrick, C. W., Batsaikhan, N., Damdinbaza, D., & Cook, J. A. (2014). Multilocus systematics and non-punctuated evolution of Holarctic Myodini (Rodentia: Arvicolinae). Molecular Phylogenetics and Evolution, 76, 18–29. doi:10.1016/j.ympev.2014.02.019

43. Kohn, M. H., & Wayne, R. K. (1997). Facts from feces revisited. Trends in Ecology & Evolution, 12(6), 223–227. doi:10.1016/S0169-5347(97)01050-1

44. Leite, E. R., & Stuth, J. W. (1994). Influence of Duration of Exposure to Field Conditions on Viability of Fecal Samples for NIRS Analysis. Journal of Range Management, 47(4), 312–314. doi:10.2307/4002553

45. Lenoir, J., & Svenning, J.-C. (2015). Climate-related range shifts – a global multidimensional synthesis and new research directions. Ecography, 38(1), 15–28. doi:10.1111/ecog.00967

46. Ley, R. E., Hamady, M., Lozupone, C., Turnbaugh, P. J., Ramey, R. R., Bircher, J. S., … Gordon, J. I. (2008). Evolution of Mammals and Their Gut Microbes. Science, 320(5883), 1647–1651. doi:10.1126/science.1155725

47. Liu, Z., Cai, W., & Shao, X. (2008). Outlier detection in near-infrared spectroscopic analysis by using Monte Carlo cross-validation. Science in China Series B: Chemistry, 51(8), 751. doi:10.1007/s11426-008-0080-x

48. Lovegrove, B. G. (2010). The allometry of rodent intestines. Journal of Comparative Physiology B, 180(5), 741–755. doi:10.1007/s00360-009-0437-2

49. Martin, M. (2011). Cutadapt removes adapter sequences from high-throughput sequencing reads. EMBnet.Journal, 17(1), 10–12. doi:10.14806/ej.17.1.200

50. Mesa-Cruz, J. B., Brown, J. L., & Kelly, M. J. (2014). Effects of natural environmental conditions on faecal glucocorticoid metabolite concentrations in jaguars (Panthera onca) in Belize. Conservation Physiology, 2(1). doi:10.1093/conphys/cou039

51. Milborrow, S. (2014). Notes on the earth package. CRAN. org. Retrieved from http://www.milbo.org/doc/earth-notes.pdf

52. Murguzur, F. J. A., Bison, M., Smis, A., Böhner, H., Struyf, E., Meire, P., & Bråthen, K. A. (2019). Towards a global arctic-alpine model for Near-infrared reflectance spectroscopy (NIRS) predictions of foliar nitrogen, phosphorus and carbon content. Scientific Reports, 9(1), 8259. doi:10.1038/s41598-019-44558-9

53. Muth, C., Oravecz, Z., & Gabry, J. (2018). User-friendly Bayesian regression modeling: A tutorial with rstanarm and shinystan. The Quantitative Methods for Psychology, 14(2), 99–119. doi:10.20982/tqmp.14.2.p099

54. Nations, J. A., & Olson, L. E. (2015). Climbing behavior of northern red-backed voles (Myodes rutilus) and scansoriality in Myodes (Rodentia, Cricetidae). Journal of Mammalogy, 96(5), 957–963. doi:10.1093/jmammal/gyv096

55. Oksanen, J., Blanchet, F. G., Friendly, M., Kindt, R., Legendre, P., McGlinn, D., … Wagner, H. (2018). vegan: Community Ecology Package. Retrieved from https://CRAN.R-project.org/package=vegan

56. Ovaskainen, O., Tikhonov, G., Norberg, A., Blanchet, F. G., Duan, L., Dunson, D., … Abrego, N. (2017). How to make more out of community data? A conceptual framework and its implementation as models and software. Ecology Letters, 20(5), 561–576. doi:10.1111/ele.12757

57. Parnell, T., Narayan, E. J., Nicolson, V., Martin-Vegue, P., Mucci, A., & Hero, J.-M. (2015). Maximizing the reliability of non-invasive endocrine sampling in the tiger ( Panthera tigris ): environmental decay and intra-sample variation in faecal glucocorticoid metabolites. Conservation Physiology, 3(1). doi:10.1093/conphys/cov053

58. Pasquini, C. (2003). Near Infrared Spectroscopy: fundamentals, practical aspects and analytical applications. Journal of the Brazilian Chemical Society, 14(2), 198–219. doi:10.1590/S0103-50532003000200006

59. Pecl, G. T., Araújo, M. B., Bell, J. D., Blanchard, J., Bonebrake, T. C., Chen, I.-C., … Williams, S. E. (2017). Biodiversity redistribution under climate change: Impacts on ecosystems and human well-being. Science, 355(6332), eaai9214. doi:10.1126/science.aai9214

60. Pettorelli, N., Laurance, W. F., O’Brien, T. G., Wegmann, M., Nagendra, H., & Turner, W. (2015). Satellite remote sensing for applied ecologists: opportunities and challenges. Journal of Applied Ecology, 839–848. doi:10.1111/1365-2664.12261@10.1111/(ISSN)1365-2664.MON_JPE

61. Ruffino, L., Oksanen, T., Hoset, K. S., Tuomi, M., Oksanen, L., Korpimäki, E., … Mäkynen, A. (2015). Predator–rodent–plant interactions along a coast–inland gradient in Fennoscandian tundra. Ecography, 39(9), 871–883. doi:10.1111/ecog.01758

62. Santos, J. P. V., Vicente, J., Villamuelas, M., Albanell, E., Serrano, E., Carvalho, J., … López-Olvera, J. R. (2014). Near infrared reflectance spectroscopy (NIRS) for predicting glucocorticoid metabolites in lyophilised and oven-dried faeces of red deer. Ecological Indicators, 45, 522–528. doi:10.1016/j.ecolind.2014.05.021

63. Saric, J., Wang, Y., Li, J., Coen, M., Utzinger, J., Marchesi, J. R., … Holmes, E. (2008). Species Variation in the Fecal Metabolome Gives Insight into Differential Gastrointestinal Function. Journal of Proteome Research, 7(1), 352–360. doi:10.1021/pr070340k

64. Schmidt, N. M., Mosbacher, J. B., Vesterinen, E. J., Roslin, T., & Michelsen, A. (2018). Limited dietary overlap amongst resident Arctic herbivores in winter: complementary insights from complementary methods. Oecologia, 187(3), 689–699. doi:10.1007/s00442-018-4147-x

65. Schwarzenberger, F. (2007). The many uses of non-invasive faecal steroid monitoring in zoo and wildlife species. International Zoo Yearbook, 41(1), 52–74. doi:10.1111/j.1748-1090.2007.00017.x

66. Sheine, W. S., & Kay, R. F. (1977). An analysis of chewed food particle size and its relationship to molar structure in the primatesCheirogaleus medius andGalago senegalensis and the insectivoranTupaia glis. American Journal of Physical Anthropology, 47(1), 15–20. doi:10.1002/ajpa.1330470106

67. Shepherd, K. D., & Walsh, M. G. (2002). Development of Reflectance Spectral Libraries for Characterization of Soil Properties. Soil Science Society of America Journal, 66(3), 988–998. doi:10.2136/sssaj2002.9880

68. Smithson, M., & Verkuilen, J. (2006). A better lemon squeezer? Maximum-likelihood regression with beta-distributed dependent variables. Psychological Methods, 11(1), 54–71. doi:10.1037/1082-989X.11.1.54

69. Soininen, E. M., Gauthier, G., Bilodeau, F., Berteaux, D., Gielly, L., Taberlet, P., … others. (2015). Highly Overlapping Winter Diet in Two Sympatric Lemming Species Revealed by DNA Metabarcoding. PloS One, 10(1), e0115335.

70. Soininen, E. M., Jensvoll, I., Killengreen, S. T., & Ims, R. A. (2015). Under the snow: a new camera trap opens the white box of subnivean ecology. Remote Sensing in Ecology and Conservation, 1(1), 29–38. doi:10.1002/rse2.2

71. Soininen, E. M., Ravolainen, V. T., Bråthen, K. A., Yoccoz, N. G., Gielly, L., & Ims, R. A. (2013). Arctic Small Rodents Have Diverse Diets and Flexible Food Selection. PLoS ONE, 8(6), e68128. doi:10.1371/journal.pone.0068128

72. Stevens, A., & Ramirez-Lopez, L. (2015). Package ‘prospectr’. Technical Report.

73. Steyaert, S. M. J. G., Hütter, F. J., Elfström, M., Zedrosser, A., Hackländer, K., Lê, M. H., … Isaksson, T. (2012). Faecal spectroscopy: a practical tool to assess diet quality in an opportunistic omnivore. Wildlife Biology, 18(4), 431–438. doi:10.2981/12-036

74. Stuth, J., Jama, A., & Tolleson, D. (2003). Direct and indirect means of predicting forage quality through near infrared reflectance spectroscopy. Field Crops Research, 84(1– 2), 45–56. doi:10.1016/S0378-4290(03)00140-0

75. Taberlet, P., Coissac, E., Pompanon, F., Gielly, L., Miquel, C., Valentini, A., … Willerslev, E. (2007). Power and limitations of the chloroplast trnL (UAA) intron for plant DNA barcoding. Nucleic Acids Research, 35(3), e14–e14. doi:10.1093/nar/gkl938

76. Taberlet, P., Gielly, L., Pautou, G., & Bouvet, J. (1991). Universal primers for amplification of three non-coding regions of chloroplast DNA. Plant Molecular Biology, 17(5), 1105–1109. doi:10.1007/BF00037152

77. Tolleson, Douglas R. (2010). Fecal NIRS: What else, what next. Shining Light on Manure Improves Livestock and Land Management, 2010, 81.

78. Tolleson, Douglas R., Randel, R. D., Stuth, J. W., & Neuendorff, D. A. (2005). Determination of sex and species in red and fallow deer by near infrared reflectance spectroscopy of the faeces. Small Ruminant Research, 57(2–3), 141–150. doi:10.1016/j.smallrumres.2004.06.020

79. Vance, C. K., Tolleson, D. R., Kinoshita, K., Rodriguez, J., & Foley, W. J. (2016). Near Infrared Spectroscopy in Wildlife and Biodiversity. Journal of Near Infrared Spectroscopy, 24(1), 1–25. doi:10.1255/jnirs.1199

80. Vesterinen, E. J., Puisto, A. I. E., Blomberg, A. S., & Lilley, T. M. (2018). Table for five, please: Dietary partitioning in boreal bats. Ecology and Evolution, 8(22), 10914– 10937. doi:10.1002/ece3.4559

81. Vesterinen, E. J., Ruokolainen, L., Wahlberg, N., Peña, C., Roslin, T., Laine, V. N., … Lilley, T. M. (2016). What you need is what you eat? Prey selection by the bat Myotis daubentonii. Molecular Ecology, 25(7), 1581–1594. doi:10.1111/mec.13564

82. Villamuelas, M., Serrano, E., Espunyes, J., Fernández, N., López-Olvera, J. R., Garel, M., … Albanell, E. (2017). Predicting herbivore faecal nitrogen using a multispecies near-infrared reflectance spectroscopy calibration. PLOS ONE, 12(4), e0176635. doi:10.1371/journal.pone.0176635

83. Villette, P., Krebs, C. J., Jung, T. S., & Boonstra, R. (2016). Can camera trapping provide accurate estimates of small mammal ( Myodes rutilus and Peromyscus maniculatus ) density in the boreal forest? Journal of Mammalogy, 97(1), 32–40. doi:10.1093/jmammal/gyv150

84. Warton, D. I., Blanchet, F. G., O’Hara, R. B., Ovaskainen, O., Taskinen, S., Walker, S. C., & Hui, F. K. C. (2015). So Many Variables: Joint Modeling in Community Ecology. Trends in Ecology & Evolution, 30(12), 766–779. doi:10.1016/j.tree.2015.09.007

85. Warton, D. I., Wright, S. T., & Wang, Y. (2012). Distance-based multivariate analyses confound location and dispersion effects. Methods in Ecology and Evolution, 3(1), 89–101. doi:10.1111/j.2041-210X.2011.00127.x

86. Whisson, D. A., Engeman, R. M., & Collins, K. (2005). Developing relative abundance techniques (RATs) for monitoring rodent populations. Wildlife Research, 32(3), 239. doi:10.1071/WR03128

87. Wickham, H. (2009). ggplot2: Elegant Graphics for Data Analysis. Springer Science & Business Media.

88. Wiedower, E. E., Kouba, A. J., Vance, C. K., Hansen, R. L., Stuth, J. W., & Tolleson, D. R. (2012). Fecal Near Infrared Spectroscopy to Discriminate Physiological Status in Giant Pandas. PLoS ONE, 7(6), e38908. doi:10.1371/journal.pone.0038908

89. Willerslev, E., Davison, J., Moora, M., Zobel, M., Coissac, E., Edwards, M. E., … Taberlet, P. (2014). Fifty thousand years of Arctic vegetation and megafaunal diet. Nature, 506(7486), 47–51. doi:10.1038/nature12921

90. Wintle, B. A., Runge, M. C., & Bekessy, S. A. (2010). Allocating monitoring effort in the face of unknown unknowns. Ecology Letters, 13(11), 1325–1337. doi:10.1111/j.1461-0248.2010.01514.x

91. Xu, Q.-S., Daszykowski, M., Walczak, B., Daeyaert, F., de Jonge, M. R., Heeres, J., … Massart, D. L. (2004). Multivariate adaptive regression splines—studies of HIV reverse transcriptase inhibitors. Chemometrics and Intelligent Laboratory Systems, 72(1), 27–34. doi:10.1016/j.chemolab.2004.02.007

92. Yoccoz, N. G. (2012). The future of environmental DNA in ecology. Molecular Ecology, 21(8), 2031–2038. doi:10.1111/j.1365-294X.2012.05505.x

93. Yoccoz, N. G., Nichols, J. D., & Boulinier, T. (2001). Monitoring of biological diversity in space and time. Trends in Ecology & Evolution, 16(8), 446–453. doi:10.1016/S0169-5347(01)02205-4

94. Zierer, J., Jackson, M. A., Kastenmüller, G., Mangino, M., Long, T., Telenti, A., … Menni, C. (2018). The fecal metabolome as a functional readout of the gut microbiome. Nature Genetics, 50(6), 790–795. doi:10.1038/s41588-018-0135-7

