## Supplemental Materials for "Novel frontier in wildlife monitoring: identification of small rodent species from faecal pellets using Near-Infrared Reflectance Spectroscopy (NIRS)"

### This supplementary material includes:

#### Detailed DNA-metabarcoring of rodent feces

After NIRS scanning, we analyzed a subset of intestinal samples from JRA and Marine site (n=385) for diet species composition with DNA-metabarcoding, where samples were divided into four sequencing runs. All wet laboratory work was performed by Center of Evolutionary Applications, University of Turku, Finland. A DNA metabarcoding approach similar to Soininen et al (2015) was used to identify plant species in vole pellets. DNA was extracted with a column based NucleoSpin Tissue kit (cat nr 740952, Macherey-Nagel, Duren, Germany) (see Soininen et al., 2015). The trnL intron in plastid DNA was amplified using universal plant primers 'g-h' and 'c-h' (for primer information see Taberlet, Gielly, Pautou, & Bouvet, 1991; Taberlet et al., 2007) and resulting fragments were sequenced on IonTorrent platform. DNA libraries were constructed using a two-step PCR approach (see e.g. Vesterinen, Puisto, Blomberg, & Lilley, 2018): the first step amplifies the target region and in the second step additional adapter and barcode sequences are added. To minimize stochastic variation in amplification, each DNA sample was replicated three times for the first PCR step after which they were pooled for the second PCR step. Amplifications from the second PCR step were purified and size selected with a SPRI bead clean-up following Vesterinen et al. (2016), concentrations were measured by Qubit (Invitrogen; www.invitrogen.com) and finally pooled in equimolar quantities. Quantity and size distribution of the pooled libraries were checked on Bioanalyzer 2100 using High Sensitivity DNA Kit (part nr 5067-4626, Agilent; www.agilent.com).

For clonal amplification of the DNA libraries we used the Ion PGM Hi-Q OT2 Kit, following the manufacturer's instructions (Life Technologies, manual cat nr 0010902, Rev A0). Template positive Ion Sphere Particles were enriched using the Ion One Touch ES. Sequencing was done on 316 v2 and 318 v2 chips with Ion PGM Hi-Q Sequencing Kit according to standard protocol, using 500 flows (Life Technologies, manual cat nr 00009816, Rev C.0). Torrent Suite 5.0.4 software was used for base calling and initial quality trimming.

#### Bioinformatics

The four sequencing runs yielded altogether 12,575,869 quality-controlled reads that could be assigned to original samples. The reads were uploaded to CSC servers (IT Center for Science, www.csc.fi) for trimming and further analysis. Trimming and quality control of the sequences were carried out partly following Schmidt et al. (2018) with several modifications, as explained below.

All the reads from the four sequencing runs were combined and trimmed for quality using 32-bit *usearch* version 11 (Edgar, 2010) with the command '*fastq_filter*' using maxee rate 1 (Edgar & Flyvbjerg, 2015). We removed primers using software *cutadapt* version 1.14 (Martin, 2011) with 20% mismatch rate and minimum trimmed length set to 8 bp following Soininen *et al.* (2013) ending with 8,351,756 trimmed sequences. Next, we collapsed sequences into unique haplotypes with command '*fastx_uniques*', denoised the unique sequences (i.e. removed chimeras) and clustered them into zero-radius OTUs (Operational Taxonomic Unit; cf. Edgar & Flyvbjerg, 2015) with command '*unoise3*' using USEARCH version 11. We mapped the OTUs back to the original trimmed reads with command '*otutab*' to establish the total number of reads in each sample using *usearch*. We were able to map 7,726,911 reads (~93% of the trimmed reads) to our original samples.

The *trnl* OTUs were initially identified to taxa using the *usearch* '*sintax*' classifier with a database consisting of artic plant trnl sequences (Willerslev et al., 2014) using 70% probability threshold for taxonomic assignation. In cases where the database was insufficient to reach even a genus-level assignation, we used local BLAST (Altschul, Gish, Miller, Myers, & Lipman, 1990) and manually assigned the taxonomy based on the knowledge of the local flora at JRA. Specifically, taxa not known to be found in the study area were assigned to their next higher taxonomic level. Similarly, taxa identified to genus level (or higher), but with only one species (or genus) present in study area, were assigned to the more detailed taxon identity. Non-regional species with closely related species found in the region were assigned to that taxon (e.g. *Cornus canadensis* to *Cornus suecica*). All observations of taxons at genus or higher level that did not occur in the region were excluded, as were observations at species level that did not have closely related species at the study area. After this, read frequencies for each diet taxon in each sample were transformed to relative read abundances (RRA; cf. Deagle et al., 2018; Vesterinen et al., 2018), as read abundances may actually be less misleading than presence/absence conversions (Deagle et al., 2018). Using the sequence counts, relative read abundance  for food item *i* is calculated as (from Deagle et al., 2018):

$$RRA_i = \frac{1}{S}\sum_{k=1}^{S}\frac{n_{i,k}}{\sum_{i=1}^{T} n_{i,k}} \times 100\% \qquad \text{(Eqn 1)}$$

where $T$ is the number of food items (taxa), $S$ is the number of samples, and $n_{i,k}$ is the number of sequences of food item $i$ in sample $k$. For subsequent analysis, we summed the RRA values at plant family level for further analysis.

#### Calibration modelling of genus and species identity

We used multivariate adaptive regression splines (MARS; Friedman, 1991), an adaptive and generalized form of stepwise linear regression (Hastie, Tibshirani, & Friedman, 2009), for calibration modelling. MARS deals well with high-dimensional input data and allows for additive effects and interactions between variables (in our case individual hinge functions of NIR-spectra; Friedman & Roosen, 1995). To test hypothesis H1, H2 and H4, we applied MARS in a flexible discriminant analysis (FDA; Hastie, Tibshirani, & Buja, 1994) framework, which fits MARS for multi-group classification problems. FDA is a generalized and restricted form of linear discriminant analysis, LDA (Hastie et al., 1994, 2009), modelling widely used in chemometrics (Pasquini, 2003). While LDA applies linear regressions on multiresponse data, FDA is a non-parametric variant and assumes non-linear decision boundaries (Hastie et al., 1994), making it suitable for the current purpose. For modelling a two-level response variable in H3, we fitted MARS directly.

In MARS and in FDA fitting MARS, model validation (i.e. estimation of prediction error for model selection, Hastie et al., 2009 p. 222) reduces the residual squared error (λ) through generalized cross validation criterion (GCV; Hastie et al., 2009 p. 244-245) of the candidate models during backward pass. This is the default procedure in MARS (Milborrow, 2014; Hastie, Tibshirani, & Buja, 2015), replacing the k-fold cross-validation commonly used in chemometric analysis due to computational reasons (cf. Pasquini, 2003; Hastie et al., 2009, Chapter 9.4).

We used Monte-Carlo cross-validation (MCCV; e.g. Xu et al., 2004) with 200 iterations within a modified repeated double cross-validation strategy (rdCV; Filzmoser, Liebmann, & Varmuza, 2009; modified, as we used GCV for variable selection) for model evaluation (hereafter *rdMCCV*). Within each iteration, we divided the modelling dataset into a joint training and validation data used for model fitting and variable selection (hereafter *calibration set*), respectively, and an *MCCV test set* for assessment of the generalization error of each iterated calibration model (reported as model misclassification rate and species prediction accuracy; Pasquini, 2003; Filzmoser et al., 2009). The main purpose of applying rdMCCV was robust hypothesis testing, i.e. to estimate how model performance and misclassification rates vary across sets of calibration data (Filzmoser et al., 2009), but also to estimate error rates for individual samples (cf. Liu, Cai, & Shao, 2008; an adaptation of the outlier detection concept ). To add robustness to the calibration models, we repeated rdMCCV procedure with 95%, 90% and 80% data splitting to calibration set, which increased the number of total rdMCCV iterations per hypothesis to 600. These splits provided the lowest misclassification rates based on preliminary comparison of ten different splits spanning from 10% to 95% of data to calibration set (data not shown). Depending on the model, the data was split as fixed percentage of each species, species and exposure treatment duration, or species, exposure treatment duration and region. We used MARS algorithms with one-way or two-way interactions of hinge functions, depending on which interaction structure provided the lowest misclassification rate of test samples based on preliminary models with 20 iterations (Filzmoser et al., 2009).

From model iterations, we extracted sample identity predictions, variable selection and variable importance data, as well as model summary statistics (including $R^2$, GCV and $R^2_{GCV}$ presented in Supplementary Table S2). Based on the sample identity predictions, we calculated the mean, standard error and 95% confidence limits for sample and taxon-specific prediction accuracy for each hypothesis and data split, as well as model misclassification rates (cf. Filzmoser et al., 2009), for all rdMCCV iterations.

#### Supplementary table S1a-d

**Supplementary Table S1a.** Composition of samples used in the study, divided by region, area within region, habitat and reproductive status. Each habitat class of JRA consists of several sub-classes, which are here pooled to heaths, mires, meadows and snowbeds. Varanger samples are a mixture of heath and meadow habitat captured individuals. Within reproductive status, visibly pregnant females are indicated separately ("female, preg"). WF = West-Finnmark, EF = East Finnmark. Species identities are coded as follows: llem = *Lemmus lemmus*, magr = *Microtus agrestis*, moec = *Microtus oeconomus*, mruf = *Myodes rufocanus* and mruti = *Myodes rutilus*.

| region | area | habitat | rep. status | llem | magr | moec | mruf | mruti |
|---|---|---|---|---|---|---|---|---|
| WF | Joatka | heath | female | 19 | 1 | | 8 | 14 |
| WF | Joatka | heath | female, preg | 4 | | | 5 | 1 |
| WF | Joatka | heath | male | 18 | | | 8 | 18 |
| WF | Joatka | mire | female | 6 | 9 | 11 | 6 | 1 |
| WF | Joatka | mire | female, preg | 2 | 4 | 2 | 11 | |
| WF | Joatka | mire | male | 18 | 21 | 10 | 10 | 2 |
| WF | Joatka | meadow | female | 3 | 10 | 41 | 7 | 31 |
| WF | Joatka | meadow | female, preg | 4 | 1 | 2 | 3 | 3 |
| WF | Joatka | meadow | male | 6 | 8 | 26 | 4 | 27 |
| WF | Joatka | meadow | NA | | | | | 1 |
| WF | Joatka | snowbed | female | 7 | | 1 | 3 | 4 |
| WF | Joatka | snowbed | female, preg | 6 | | | 1 | |
| WF | Joatka | snowbed | male | 14 | 3 | | 3 | 1 |
| WF | Marine | heath | female | | 2 | | 1 | |
| WF | Marine | heath | male | | 10 | | 1 | |
| WF | Marine | mire | female | | 1 | | 1 | |
| WF | Marine | mire | female, preg | | 1 | | | |
| WF | Marine | mire | male | | 3 | | | |
| WF | Marine | snowbed | female | | 10 | | | |
| WF | Marine | snowbed | female, preg | | 3 | | | |
| WF | Marine | snowbed | male | | 10 | | | |
| WF | total | | | 107 | 97 | 93 | 72 | 103 |
| EF | Ifjord | | female | 1 | | 17 | 15 | |
| EF | Ifjord | | female, preg | | | 3 | | |
| EF | Ifjord | | male | | | 8 | 9 | |
| EF | Ifjord | | NA | | | | 1 | |
| EF | Komagdalen | | female | | | 5 | | |
| EF | Komagdalen | | male | | | 9 | | |
| EF | Vjaokbselv | | female | | | | 7 | 1 |
| EF | Vjaokbselv | | female, preg | | | | 1 | |
| EF | Vjaokbselv | | male | 1 | | | 5 | |
| EF | total | | | 2 | 0 | 42 | 38 | 1 |

**Supplementary Table S1b.** Number of JRA samples in 5%, 10% and 20% data splits used for H1 and H2 calibration models divided by duration of exposure treatment (weeks) and species. Species identities are coded as follows: llem = *Lemmus lemmus*, magr = *Microtus agrestis*, moec = *Microtus oeconomus*, mruf = *Myodes rufocanus* and mruti = *Myodes rutilus*.

| week | species | 5% | 10% | 20% |
|---|---|---|---|---|
| 0 | llem | 102 | 96 | 86 |
| 0 | magr | 92 | 87 | 78 |
| 0 | moec | 88 | 84 | 74 |
| 0 | mruf | 68 | 65 | 58 |
| 0 | mruti | 98 | 93 | 82 |
| 1 | llem | 48 | 45 | 40 |
| 1 | magr | 24 | 23 | 20 |
| 1 | moec | 24 | 23 | 20 |
| 1 | mruf | 47 | 44 | 39 |
| 2 | llem | 48 | 45 | 40 |
| 2 | magr | 24 | 23 | 20 |
| 2 | moec | 24 | 23 | 20 |
| 2 | mruf | 48 | 45 | 40 |
| 3 | llem | 48 | 45 | 40 |
| 3 | magr | 24 | 23 | 20 |
| 3 | moec | 23 | 22 | 19 |
| 3 | mruf | 44 | 41 | 37 |
| 4 | llem | 46 | 43 | 38 |
| 4 | magr | 24 | 23 | 20 |
| 4 | moec | 24 | 23 | 20 |
| 4 | mruf | 48 | 45 | 40 |
| 5 | llem | 33 | 31 | 28 |
| 5 | magr | 24 | 23 | 20 |
| 5 | moec | 24 | 23 | 20 |
| 5 | mruf | 48 | 45 | 40 |
| 6 | llem | 24 | 23 | 20 |
| 6 | magr | 24 | 23 | 20 |
| 6 | moec | 10 | 10 | 9 |
| 6 | mruf | 43 | 41 | 36 |

**Supplementary Table S1c.** Number of samples in 5%, 10% and 20% data splits used for H3 calibration models. Species identities are coded as follows: moec = *Microtus oeconomus* and mruf = *Myodes rufocanus.*

| **hypothesis** | **species** | **region** | **5%** | **10%** | **20%** |
|---|---|---|---|---|---|
| H3.1 | moec | JRA | 217 | 208 | 182 |
| H3.1 | mruf | JRA | 346 | 326 | 290 |
| H3.2 | moec | Varanger | 40 | 38 | 34 |
| H3.2 | mruf | Varanger | 36 | 34 | 30 |
| H3.3 | moec | both | 257 | 246 | 216 |
| H3.3 | mruf | both | 382 | 360 | 320 |

**Supplementary Table S1d.** Number of WF week 0 samples in 5%, 10% and 20% data splits used for H4 calibration models. Species identities are coded as follows: llem = *Lemmus lemmus*, magr = *Microtus agrestis*, moec = *Microtus oeconomus*, mruf = *Myodes rufocanus* and mruti = *Myodes rutilus*.

| species | 5% | 10% | 20% |
|---|---|---|---|
| llem | 94 | 89 | 79 |
| magr | 86 | 81 | 72 |
| moec | 80 | 76 | 67 |
| mruf | 62 | 59 | 52 |
| mruti | 40 | 38 | 34 |

#### Supplementary Table S2a-b

**Supplementary Table S2a.** Model summary statistics for H1 and H2 species and genus models. Table includes the mean, standard deviation, minimum and maximum of $R^2$, GCV and $R^2_{GCV}$ values across 200 iterations per each data split.

| statistic | hypothesis | response | data_split | mean | sd | min | max |
|---|---|---|---|---|---|---|---|
| $R^2$ | H1 | genus | 10% | 0.852381 | 0.024222 | 0.782172 | 0.900066 |
| | | | 20% | 0.853877 | 0.022328 | 0.786243 | 0.901558 |
| | | | 5% | 0.852657 | 0.024314 | 0.778218 | 0.899053 |
| | | species | 10% | 0.723132 | 0.030292 | 0.631264 | 0.788646 |
| | | | 20% | 0.725762 | 0.034838 | 0.618169 | 0.814711 |
| | | | 5% | 0.720956 | 0.03014 | 0.643815 | 0.822423 |
| | H2 | genus | 10% | 0.936394 | 0.006219 | 0.897083 | 0.951105 |
| | | | 20% | 0.939206 | 0.004728 | 0.9191 | 0.949427 |
| | | | 5% | 0.934538 | 0.006522 | 0.904935 | 0.948589 |
| | | species | 10% | 0.888679 | 0.009632 | 0.845349 | 0.908743 |
| | | | 20% | 0.88613 | 0.011633 | 0.852072 | 0.908929 |
| | | | 5% | 0.887466 | 0.009004 | 0.845703 | 0.909189 |
| GCV | H1 | genus | 10% | 0.380685 | 0.037559 | 0.307389 | 0.497703 |
| | | | 20% | 0.387012 | 0.034424 | 0.322674 | 0.489662 |
| | | | 5% | 0.376432 | 0.038074 | 0.307268 | 0.496674 |
| | | species | 10% | 1.410264 | 0.074131 | 1.252336 | 1.669844 |
| | | | 20% | 1.435192 | 0.083356 | 1.259031 | 1.731559 |
| | | | 5% | 1.407058 | 0.072597 | 1.184209 | 1.615848 |
| | H2 | genus | 10% | 0.215609 | 0.011339 | 0.191873 | 0.278563 |
| | | | 20% | 0.220395 | 0.010539 | 0.188329 | 0.257919 |
| | | | 5% | 0.214543 | 0.010838 | 0.195404 | 0.258646 |
| | | species | 10% | 0.711401 | 0.025487 | 0.653289 | 0.77878 |
| | | | 20% | 0.763039 | 0.035775 | 0.671267 | 0.898398 |
| | | | 5% | 0.69878 | 0.024462 | 0.623395 | 0.787014 |
| $R^2_{GCV}$ | H1 | genus | 10% | 0.80998 | 0.018747 | 0.75157 | 0.846566 |
| | | | 20% | 0.806864 | 0.017179 | 0.755638 | 0.838972 |
| | | | 5% | 0.812086 | 0.019007 | 0.752062 | 0.846613 |
| | | species | 10% | 0.648031 | 0.018501 | 0.583246 | 0.687446 |
| | | | 20% | 0.641889 | 0.020799 | 0.567939 | 0.685845 |
| | | | 5% | 0.6488 | 0.01812 | 0.596686 | 0.704423 |
| | H2 | genus | 10% | 0.892481 | 0.005655 | 0.861087 | 0.904317 |
| | | | 20% | 0.890133 | 0.005254 | 0.871427 | 0.906118 |
| | | | 5% | 0.892997 | 0.005406 | 0.871001 | 0.902543 |
| | | species | 10% | 0.763494 | 0.008473 | 0.741094 | 0.782814 |
| | | | 20% | 0.746417 | 0.011889 | 0.701433 | 0.776916 |
| | | | 5% | 0.767657 | 0.008134 | 0.738319 | 0.792722 |

**Supplementary Table S2b.** Model summary statistics for H4 models, more specifically $R^2$, GCV and $R^2_{GCV}$ values. Table includes the mean, standard deviation and minimum and maximum values across 200 iterations per each data split.

| statistic | data_split | mean | sd | min | max |
|---|---|---|---|---|---|
| Rsq | 10% | 0.702589 | 0.041305 | 0.584969 | 0.802068 |
| Rsq | 20% | 0.689066 | 0.041683 | 0.567168 | 0.794938 |
| Rsq | 5% | 0.703459 | 0.03706 | 0.582963 | 0.785231 |
| GCV | 10% | 1.971175 | 0.076891 | 1.780157 | 2.227043 |
| GCV | 20% | 2.031704 | 0.095246 | 1.720192 | 2.35227 |
| GCV | 5% | 1.945534 | 0.071902 | 1.718549 | 2.121494 |
| GCV_Rsq | 10% | 0.510075 | 0.019111 | 0.446481 | 0.557552 |
| GCV_Rsq | 20% | 0.49541 | 0.023655 | 0.415795 | 0.572777 |
| GCV_Rsq | 5% | 0.5163 | 0.017876 | 0.472553 | 0.572733 |

### Supplementary Table S3

**Supplementary Table S3.** Posterior distributions of model parameters. Independent variable (week) distributions not including zero are denoted with bold, and 95% confidence interval with bold italics.

| Parameter | hypothesis | species | mean | se | sd | 2.5% CI | 97.5% CI | n_eff | Rhat |
|---|---|---|---|---|---|---|---|---|---|
| Intercept | H1 | L. lemmus | 2.469 | 0.003 | 0.120 | 2.226 | 2.701 | 2104 | 1.001 |
| | H1 | M. agrestis | 0.543 | 0.002 | 0.122 | 0.302 | 0.785 | 3264 | 1.001 |
| | H1 | M. oeconomus | 0.364 | 0.002 | 0.125 | 0.128 | 0.615 | 3538 | 1.000 |
| | H1 | M. rufocanus | 2.110 | 0.003 | 0.129 | 1.847 | 2.361 | 1791 | 1.000 |
| | H2 | L. lemmus | 3.120 | 0.004 | 0.180 | 2.768 | 3.473 | 1750 | 1.001 |
| | H2 | M. agrestis | 2.015 | 0.005 | 0.227 | 1.582 | 2.462 | 1949 | 1.002 |
| | H2 | M. oeconomus | 1.270 | 0.005 | 0.251 | 0.783 | 1.750 | 2557 | 1.000 |
| | H2 | M. rufocanus | 2.840 | 0.004 | 0.172 | 2.508 | 3.182 | 2001 | 1.000 |
| Phi | H1 | L. lemmus | 2.033 | 0.005 | 0.217 | 1.634 | 2.477 | 1991 | 1.001 |
| | H1 | M. agrestis | 0.551 | 0.001 | 0.042 | 0.471 | 0.636 | 2574 | 1.003 |
| | H1 | M. oeconomus | 0.557 | 0.001 | 0.043 | 0.479 | 0.643 | 3085 | 1.000 |
| | H1 | M. rufocanus | 1.508 | 0.004 | 0.152 | 1.221 | 1.814 | 1684 | 1.001 |
| | H2 | L. lemmus | 4.413 | 0.015 | 0.603 | 3.329 | 5.656 | 1548 | 1.001 |
| | H2 | M. agrestis | 2.632 | 0.009 | 0.375 | 1.961 | 3.422 | 1859 | 1.003 |
| | H2 | M. oeconomus | 1.632 | 0.005 | 0.220 | 1.239 | 2.109 | 2108 | 1.001 |
| | H2 | M. rufocanus | 4.358 | 0.013 | 0.541 | 3.370 | 5.465 | 1715 | 1.002 |
| log-posterior | H1 | L. lemmus | 1617.613 | 0.029 | 1.203 | 1614.465 | 1618.995 | 1673 | 1.003 |
| | H1 | M. agrestis | 587.172 | 0.028 | 1.239 | 583.891 | 588.581 | 2003 | 1.001 |
| | H1 | M. oeconomus | 421.345 | 0.027 | 1.228 | 418.334 | 422.754 | 2035 | 1.000 |
| | H1 | M. rufocanus | 1486.889 | 0.035 | 1.269 | 1483.429 | 1488.289 | 1345 | 1.001 |
| | H2 | L. lemmus | 1102.104 | 0.030 | 1.199 | 1098.972 | 1103.451 | 1636 | 1.001 |
| | H2 | M. agrestis | 312.183 | 0.032 | 1.287 | 308.776 | 313.595 | 1576 | 1.000 |
| | H2 | M. oeconomus | 230.857 | 0.030 | 1.227 | 227.679 | 232.265 | 1640 | 1.002 |
| | H2 | M. rufocanus | 1058.271 | 0.033 | 1.279 | 1054.925 | 1059.694 | 1501 | 1.003 |
| mean_PPD | H1 | L. lemmus | 0.925 | 0.000 | 0.011 | 0.902 | 0.944 | 2601 | 1.000 |
| | H1 | M. agrestis | 0.688 | 0.001 | 0.031 | 0.624 | 0.748 | 3391 | 1.000 |
| | H1 | M. oeconomus | 0.646 | 0.001 | 0.033 | 0.581 | 0.708 | 3930 | 1.000 |
| | H1 | M. rufocanus | 0.899 | 0.000 | 0.014 | 0.871 | 0.925 | 2447 | 1.000 |
| | H2 | L. lemmus | 0.958 | 0.000 | 0.008 | 0.941 | 0.971 | 2284 | 1.000 |
| | H2 | M. agrestis | 0.893 | 0.000 | 0.018 | 0.855 | 0.925 | 2603 | 1.001 |
| | H2 | M. oeconomus | 0.834 | 0.001 | 0.028 | 0.775 | 0.884 | 2922 | 1.000 |
| | H2 | M. rufocanus | 0.951 | 0.000 | 0.008 | 0.935 | 0.965 | 2397 | 1.000 |
| week | H1 | L. lemmus | 0.022 | 0.001 | 0.029 | -0.033 | 0.078 | 2276 | 1.000 |
| | **H1** | **M. agrestis** | **0.124** | **0.001** | **0.039** | ***0.049*** | ***0.199*** | **3072** | **1.000** |
| | **H1** | **M. oeconomus** | **0.131** | **0.001** | **0.044** | ***0.046*** | ***0.216*** | **3596** | **0.999** |
| | H1 | M. rufocanus | 0.032 | 0.001 | 0.028 | -0.023 | 0.088 | 2209 | 1.001 |
| | H2 | L. lemmus | 0.004 | 0.001 | 0.040 | -0.075 | 0.079 | 2230 | 1.000 |
| | H2 | M. agrestis | 0.034 | 0.001 | 0.052 | -0.070 | 0.139 | 2567 | 1.000 |
| | H2 | M. oeconomus | 0.113 | 0.001 | 0.067 | -0.017 | 0.244 | 2625 | 1.000 |
| | H2 | M. rufocanus | 0.039 | 0.001 | 0.038 | -0.036 | 0.112 | 1998 | 1.001 |

#### Supplementary Figure S1

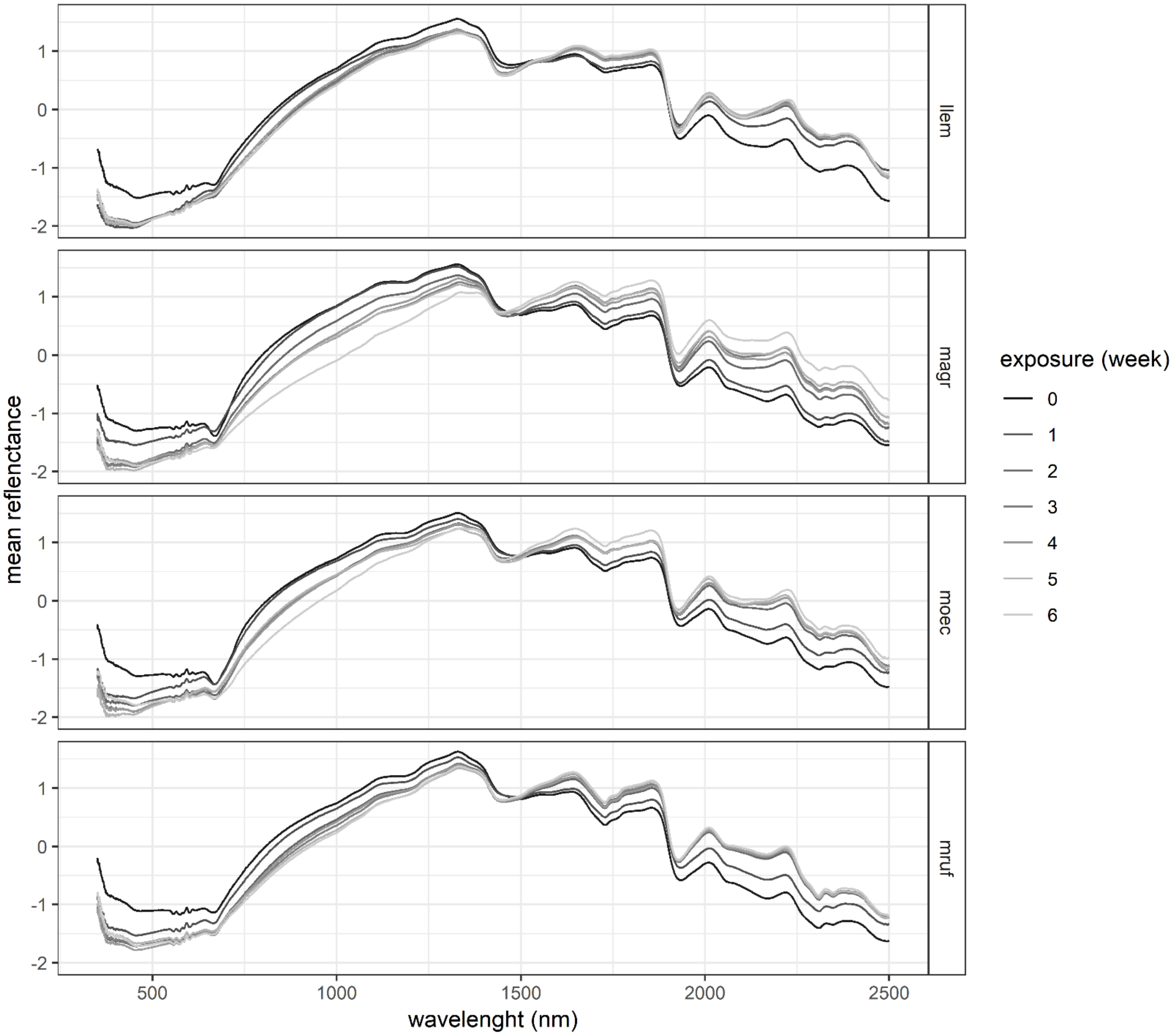

**Supplementary Figure S1**. Mean reflectance spectra of four species included in the exposure treatment. Spectra of intestinal samples and each exposure week are plotted separately, with lighter grey indicating increasing time of exposure. Species names: llem = *Lemmus lemmus*, magr = *Microtus agrestis*, moec = *Microtus oeconomus*, mruf = *Myodes rufocanus*.

### Supplementary Figure S2

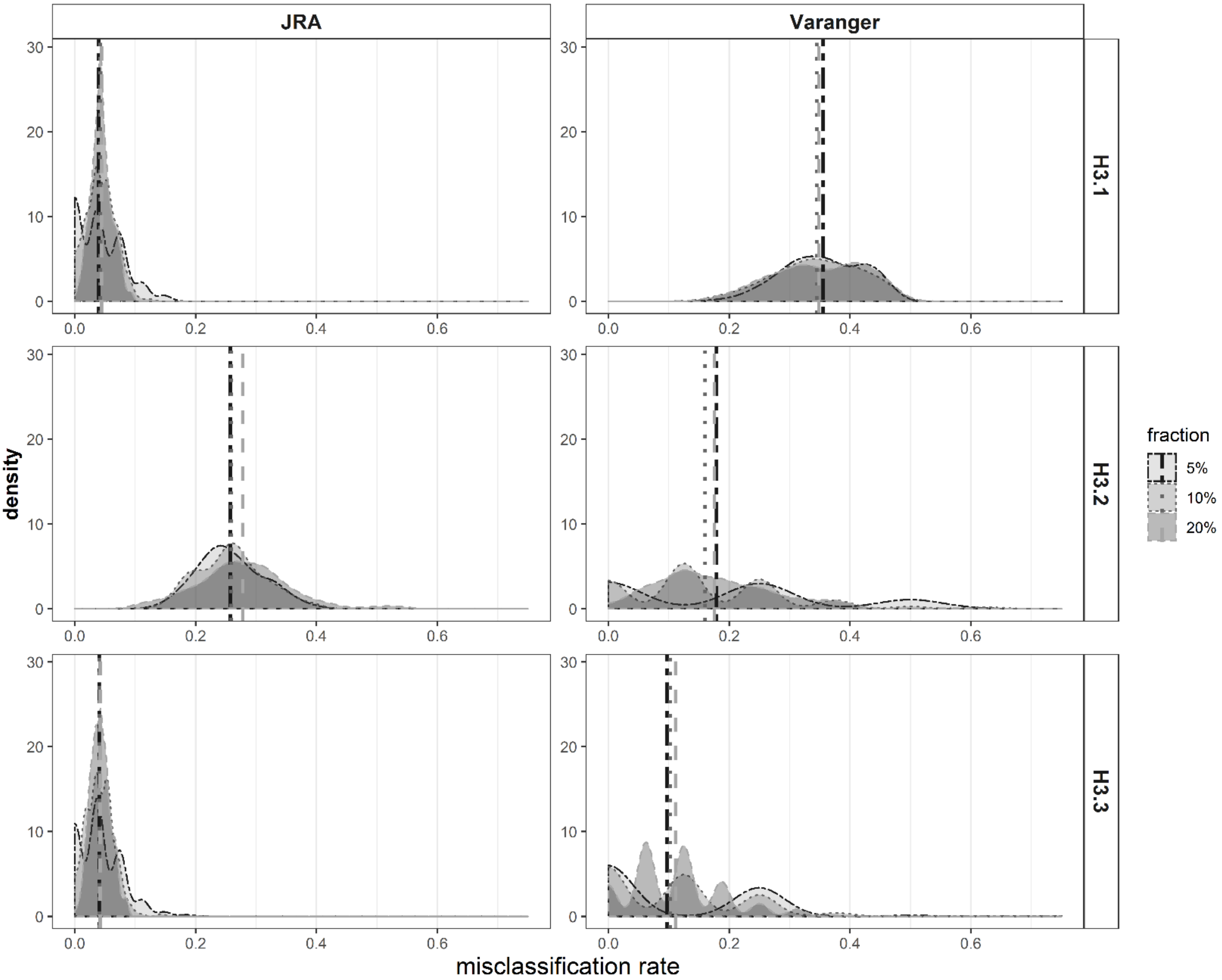

**Supplementary Figure S2.** Density plots of misclassification rate of MCCV test set samples in H3 models. Model iterations (n=200) with different data splits (fractions) are plotted separately.

#### Supplementary Figure S3

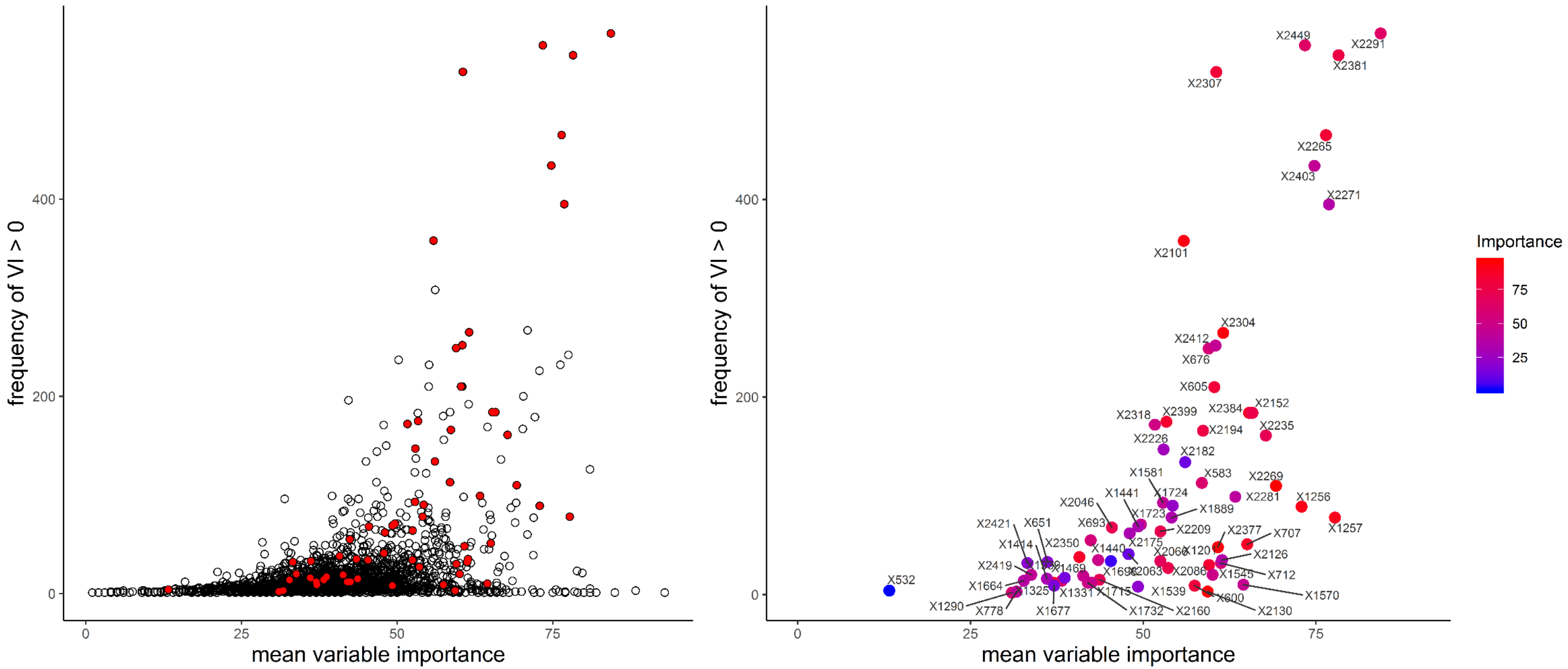

**Supplementary Figure S4.** Mean variable importance and frequency of VI > 0 across 600 MCCV runs with TOC modeling data. Left panel: variables selected in TOC model plotted with red fill, while variables selected across 600 MCCV runs but not in the TOC model are plotted with transparent fill. Right panel: Only variables selected in the TOC model plotted, with color indicating variable importance, and position in xy-space the MCCV model variable selection.
